# Joint contributions of cortical morphometry and white matter microstructure in healthy brain aging: A partial least squares correlation analysis

**DOI:** 10.1101/620419

**Authors:** David A. Hoagey, Jenny R. Rieck, Karen M. Rodrigue, Kristen M. Kennedy

## Abstract

Cortical atrophy and degraded axonal health have been shown to coincide during normal aging; however, few studies have examined these measures together. To lend insight into both the regional specificity and the relative timecourse of structural degradation of these tissue compartments across the lifespan, we analyzed grey matter (GM) morphometry (cortical thickness, surface area, volume) and estimates of white matter (WM) microstructure (fractional anisotropy, mean diffusivity) using traditional univariate and more robust multivariate techniques to examine age associations in 186 healthy adults aged 20-94 years old. Univariate analysis of each tissue type revealed that negative age associations were largest in frontal grey and white matter tissue and weaker in temporal, cingulate, and occipital regions, representative of not only an anterior-to-posterior gradient, but also a medial-to-lateral gradient. Multivariate partial least squares correlation (PLSC) found the greatest covariance between GM and WM was driven by the relationship between WM metrics in the anterior corpus callosum and projections of the genu, anterior cingulum, and fornix; and with GM thickness in parietal and frontal regions. Surface area was far less susceptible to age effects and displayed less covariance with WM metrics, while regional volume covariance patterns largely mirrored those of cortical thickness. Results support a retrogenesis-like model of aging, revealing a coupled relationship between frontal and parietal GM and the underlying WM, which evidence the most protracted development and the most vulnerability during healthy aging.

## Introduction

The healthy adult brain is susceptible to structural degradation of both grey and white matter tissue throughout the aging process. To understand the relative timecourse and anatomical specificity of volumetric decline, much effort has been directed toward associating loss of structural integrity, as observed through various magnetic resonance imaging (MRI) modalities, with increasing age. The majority of these aging studies utilized a univariate approach to assess grey matter and white matter tissue compartments separately for their sensitivity to the effects of time. Cumulatively, these studies demonstrated the differential age-sensitivity of regional grey matter and white matter properties, but leave unclear how these tissue compartments influence each other and age together. Given that these structural measurements reflect components of one cellular unit – the neuron, with somatic, dendritic, and axonal components, it stands to reason that aging of these biological components must be linked. However, few studies have taken this dependency into account by examining the joint contribution of both grey and white matter tissue to investigate how alterations in both aspects of brain structure are related across the lifespan. In contrast to univariate approaches, multivariate analysis techniques are perfectly poised to address this issue. As demonstrated by Groves et al., 2012, information from multiple imaging modalities can be analyzed together to assess the combined effect of structural integrity loss using multivariate analysis techniques. Treating complementary components of the neuronal unit as a whole, as opposed to independent features with orthogonal contributions, allows for a more complete representation of imaging data. Multimodal analyses such as these, which synthesize interdependent data, are necessary to advance understanding of the progression of structural decline and the influence on cognition throughout the lifespan.

Advances in MRI pulse sequences, and subsequent imaging analysis processing tools, allowed for an abundance of research identifying age-related associations in both cortical morphometry and white matter health. Regional differences in cortical morphometry associated with age (Raz, 2000; Raz and Rodrigue, 2006) generally show measurement peaks during childhood (Tamnes, et al., 2009), giving way to an anatomically and chronologically divergent pattern of decline during senescence (Fjell, et al., 2013; Shaw, et al., 2008). Regional analyses revealed age-related atrophy in frontal (Allen, et al., 2005; Bartzokis, et al., 2001; Fjell, et al., 2009; Grieve, et al., 2005; Jernigan, et al., 2001; Lemaitre, et al., 2012; Manard, et al., 2016; Resnick, et al., 2003; Sowell, et al., 2003; Zimmerman, et al., 2006), parietal (Fjell, et al., 2009; Grieve, et al., 2005; Sowell, et al., 2003), and select temporal (Bartzokis, et al., 2001; Fjell, et al., 2009; Sowell, et al., 2003) regions of grey matter, yet relatively no association with limbic cortices (Grieve, et al., 2005). Biological mechanisms underlying regional vulnerability to aging include loss of synaptic density and/or dendritic arborization (Raz, 2001; Salat, et al., 2004), alterations in small diameter or thinly myelinated intracortical white matter (Eickhoff, et al., 2005; Walters, et al., 2003; Westlye, et al., 2010; Ziegler, et al., 2010), neuronal shrinkage (Sowell, et al., 2004), and/or loss of neuropil (Lemaitre, et al., 2012; Pakkenberg, et al., 2003; Ziegler, et al., 2010). Measures of white matter health across the lifespan often demonstrate an inverted U-shaped curve with peak levels of estimated integrity during early adulthood (Raz, 2000; Westlye, et al., 2009). After a short plateau period, the fourth or fifth decade of life initiates an acceleration of age-related structural loss, which continues throughout senescence (for review see Bennett and Madden, 2014).

Macroscale aspects of cortical morphometry have been shown to represent unique contributions to age-related atrophy, for example age-related differences in volume appear to be differentially driven by its two components, cortical thickness and cortical surface area with surface area remaining relatively age-invariant and cortical thickness demonstrating particular vulnerability to the aging process (Lemaitre, et al., 2012; Panizzon, et al., 2012). Differences observed among these measures are likely to be driven by cyto- and myelo-architectonic differences across cortical tissue and the way in which different estimates account for the ratio of white and grey matter within regionally distinct laminar structure (Lemaitre, et al., 2012; Paus, et al., 2008; Raz, 2001; Storsve, et al., 2016).

Quantification of regional specificity in age-related structural brain decline prompted the development of different theories of brain aging to explain the reported patterns of cross-sectional age-related differences and longitudinal decline (Salat, 2011). For example, typically, white matter of the frontal lobes shows greater loss of integrity than do more posterior regions (Bennett, et al., 2010; Kennedy and Raz, 2009; Madden, et al., 2009; Salat, et al., 2005). This gradient-like relationship has been observed throughout the white matter of the cortex and has been described as a frontal vulnerability, or as an anterior-to-posterior gradient (Davis, et al., 2009; Head, et al., 2004; Pfefferbaum, et al., 2005; Ziegler, et al., 2010). Other directional age-gradients may exist, including increased age-vulnerability in superior white matter compared to inferior and greater vulnerability in lateral vs medial white matter (Sexton, et al., 2014; Sullivan, et al., 2010). In addition, regionally specific structural losses in aging have been proposed to mirror evolutionary and developmental trajectories through a retrogenesis or “last-in, first-out” framework of decline (Raz, 2000). Many of the later myelinating association fibers that connect higher-order cognitive association centers, exhibit greater age-related integrity loss than fibers in earlier developing limbic or primary visual, sensory, or motor areas (Salat, et al., 2005; Ziegler, et al., 2010).

Although MRI measures serve only as a proxy for the underlying cellular architecture, it seems apparent that these measured MRI signals track alterations to the neuronal components. Given the intimate connection among cell body, axons, and dendrites, aging should lead to non-independent alterations to these MRI-based grey and white matter metrics (Gao, et al., 2017; Groves, et al., 2012; Salat, et al., 2004; Storsve, et al., 2016). Specificity in this coupled aging include proposed lead-lag relationships such that cortical thinning via cell body damage leads to axonal alteration and decreased white matter health (e.g., Wallerian degeneration), or that decreased axonal health propagates from the axon to produce alterations in the cell body (e.g., transneuronal atrophy) (Kochunov, et al., 2011; Storsve, et al., 2016). Given this grey matter-white matter dependency, studies have begun to incorporate the idea of coupled age-differences (Brickman, et al., 2007) and coupled decline (Brickman, et al., 2007; Raz, et al., 2005). Univariate correlational analyses revealed associations between aging trajectories of cortical morphometry and indices of white matter health (such as fractional anisotropy (FA), which is sensitive to the directionality of water flow, or mean diffusivity (MD), which measures the overall rate of diffusion; (Kochunov, et al., 2011; Storsve, et al., 2016) and the underlying intracortical myelin (He, et al., 2007; Vidal-Pineiro, et al., 2016). Additionally, multivariate frameworks can be utilized to make accurate predictions of age when accounting for age-related patterns of covariance among multiple tissue types (Groves, et al., 2012). While current *in-vivo* human neuroimaging techniques are not yet capable of fine-grained analyses at the neuronal level, improved statistical techniques, which properly account for the relatedness of these neuronal proxies, can and should now be employed to examine the coupled relationship of neuronal component data. This approach could shed light on regional differences in how neuronal components age together across the cortex and will bridge gaps in our knowledge of how this structural covariance varies with aging.

The current study aims to first utilize univariate analyses of each tissue type in an attempt to replicate and enhance previous lifespan aging work, examining linear and quadratic age associations with grey matter cortical thickness, volume, and surface area, as well as with white matter fractional anisotropy and mean diffusivity; and second to utilize multivariate analyses (partial least squares correlation; PLSC) combining, in turn, measures of grey matter thickness, surface area, and volume, with white matter fractional anisotropy and mean diffusivity to evaluate covariance patterns. We then consider the results from these approaches in the context of two major theoretical models of brain aging: the last-in first-out retrogenesis pattern and directional/spatial age-gradients patterns (e.g., anterior-to-posterior). We predict that univariate analyses will demonstrate a last-in first-out trend where the association cortices, and separately, their underlying connections, are associated with the strongest age differences as opposed to primary and secondary sensory cortex. Further, we predict that a multivariate analysis will expand on these findings and reveal regionally differential coupled grey matter-white matter associations, such that higher order association cortices and their underlying white matter fibers, will show the greatest coupling of structural age-differences.

## Materials and Methods

### Participants

Participants included 190 cognitively normal healthy adults sampled across the lifespan ranging in age from 20-94 years of age (mean age = 53.7 years; 110 women) recruited by flyers and media ads from the Dallas-Fort Worth Metroplex. Participants were screened to be free from a history of neurological, cardiovascular, metabolic, or psychiatric problems, head trauma involving loss of consciousness, substance abuse, or cognitive altering medications. Participants were also free from MRI contraindications, such as metallic implants and claustrophobia. Further inclusion criteria required Mini-Mental State Exam (MMSE) scores > 25 (Folstein, et al., 1975) and Center for Epidemiologic Studies Depression Scale (CES-D) scores < 16 (Radloff, 1977). Before entering the study, each participant provided written informed consent, in accord with the local Institutional Review Boards. Participant demographic data are summarized in Table 1, broken down by arbitrary age groups. In total, four participants were excluded from data analysis: low MMSE score (*n* = 1), incorrect neuroimaging data acquisition (*n* = 1), abnormalities in brain structure (*n* = 2), yielding a total *N* = 186.

**Table 1.** Sample demographics.

| **Age Group**<br>**(range)** | **Sample Size**<br>**(M/F)** | **Age**<br>**mean ± SD** | **Education**<br>**mean ± SD** | **MMSE**<br>**mean ± SD** |
| --- | --- | --- | --- | --- |
| YA (20-34) | 43 (19/24) | 27.58 ± 4.37 | 15.63 ± 2.12 | 29.16 ± 0.94 |
| MA (35-54) | 46 (21/25) | 45.72 ± 5.36 | 15.26 ± 2.51 | 29.28 ± 0.80 |
| OA (55-69) | 48 (17/31) | 61.23 ± 3.66 | 15.58 ± 2.34 | 28.85 ± 0.82 |
| VOA (70-94) | 49 (19/30) | 76.75 ± 5.87 | 15.65 ± 2.77 | 28.80 ± 0.70 |
| **Total (20-94)** | **186 (76/110)** | **53.70 ± 18.77** | **15.53 ± 2.46** | **29.02 ± 0.70** |

Note. YA = Younger Adults; MA = Middle-Aged Adults; OA = Older Adults; VOA = Very Old Adults; M = Male; F = Female; Education = years of education; MMSE = Mini-Mental State Exam; SD = Standard Deviation. Age groups are arbitrarily provided to illustrate sample distribution; however, all analyses used age as a continuous variable.

### MRI Protocol

All participants were scanned on the same 3-Tesla Philips Achieva scanner with a 32-channel head coil using SENSE encoding (Philips Healthcare Systems, Best, Netherlands). A diffusion weighted imaging single shot EPI sequence was acquired with the following parameters: 65 axial slices with voxel size of 2×2×2.2 mm^3^ (reconstructed to .85×.85×2.2mm^3^), 30 diffusion weighted directions (b-value = 1000s/mm^2^) with 1 non-diffusion weighted b_0_ (0 s/ mm^2^), TR/TE = 5608ms/51ms, FOV = 224×224, matrix = 112×112, 4:19min. A high-resolution T1-weighted MPRAGE sequence was acquired with the following parameters: 160 sagittal slices, voxel size 1×1×1mm^3^, flip angle = 12°, TR/TE/TI = 8.1ms/3.7ms/1100ms, FOV = 256×204×160, matrix = 256×256, 3:57min. T2 FLAIR images were collected using the following parameters: 64 axial slices, voxel size .5×.5×2.5 mm^3^, flip angle=90°, TR/TE/TI = 11000ms/125ms/2800ms, FOV=230×230, matrix=512×512, 3:40min.

### MRI Data Processing

After visual inspection for subject movement distortions and acquisition artifacts, cortical reconstruction and volumetric segmentation was completed using Freesurfer v5.3 image analysis suite (Dale, et al., 1999; Fischl and Dale, 2000). Freesurfer includes motion correction and tissue segmentation procedures allowing isolation and quantification of tissue properties such as volume, thickness, and surface area. Furthermore, Freesurfer includes tools to segment regions of interest (ROI) which allows for alignment of a pre-defined atlas to the anatomy of an individual’s T1 image. Dura removal and control point edits were manually performed by trained researchers to optimize results and reliability. A standard atlas of cortical parcellation (Desikan, et al., 2006) was used to extract the morphometry measures (cortical thickness, surface area, volume) from 34 distinct parcellations in each hemisphere for all participants.

Diffusion images were preprocessed using the DTIPrep v1.2.4 quality control software suite to detect acquisition artifacts including susceptibility, eddy current, and subject movement distortions (Liu, et al., 2010). Using the default settings, slice-wise and gradient-wise artifacts, appearing as intensity distortions, were corrected by removing associated gradients from analysis. On average, less than four gradients were removed per subject. Appropriate corrections were applied to minimize the effects of distortions, including those caused by head motion in the scanner, by removing gradients determined to be of insufficient quality, at the default threshold levels, and by registering all remaining gradients to the non-weighted b_0_ image. Diffusion directions were adjusted to account for independent rotations of any gradient relative to the original encoding direction (Leemans and Jones, 2009). Diffusion tensors were calculated using the DSI Studio software package build from September 26^th^, 2014 (Yeh, et al., 2013). Diffusion metrics of fractional anisotropy (FA) and mean diffusivity (MD) were calculated at each voxel.

Finally, a study-specific template was created from all FA images to align all voxels using Advanced Normalization Tools (ANTs) template creation and image registration tools (Avants, et al., 2009). This template aligned all voxels within all participants to allow for analyses to be spatially equivalent. To further refine our analyses to voxels which are equivalent across the sample we restricted the template to only contain voxels with FA values > .15 for all participants. Any voxel with a value below .15 was not considered to be white matter and was removed from all analyses for all participants. The final WM space contained 138,283 voxels. A similar procedure was applied to the whole brain MD images, but we used the same voxels derived from the refinement procedure described above to create an identical template.

The influence of white matter hyperintensities (WMH), and any potential bias they could introduce to our analyses, was of concern given previous research (Jones, et al., 1999; Vernooij, et al., 2008). In our sample of healthy adults, 108 participants were identified as having WMH. The median participant from this subsample only had WMH in 0.39% (541 voxels) of the white matter voxels in the template. Despite this low number, we ran all analyses twice; once with WMH included, and once with WMH excluded in such a way that any voxel containing a WMH for any participant was removed for all participants to ensure that every participant had data at every voxel. There were no differences between the two methods and thus no bias of WMH under this analysis method. Given the ultra-conservative nature of removing any WMH voxel from all participants and the discontinuity it creates in our WM template; we present the data which include WMH voxels.

### Data Analysis Procedures

#### Univariate Analyses

Separate univariate analyses were conducted to estimate the effect of age on both grey and white matter tissue. The three grey matter morphometry measures (thickness, volume and surface area) were extracted for each ROI and averaged across hemispheres within each individual. Statistical models were conducted using R statistical software (R Core Team, 2016) within RStudio (RStudio Team, 2016) to estimate the linear and quadratic effects of age (as continuous variables) on regional cortical thickness, surface area, and volume, covarying for sex. To test for statistical significance, permutation resampling without replacement was used to build a null distribution of *t*-values for each region. Non-parametric *p*-values were calculated by dividing the number of permuted *t*-values that were less than the observed *t*-value by the total number of permutations conducted (i.e. permuted < observed / 10,000). White matter voxels were analyzed using FSL’s permutation-based general linear model program randomise (Winkler, et al., 2014) to test the linear and quadratic effect of age (as a continuous variable) on FA and MD, covarying for sex. Family-wise error corrected *p*-values were calculated at each voxel after 10,000 (Dickie, et al., 2015) permutations were conducted with Threshold-Free Cluster Enhancement.

#### Multivariate Analyses

Data were combined and analyzed using the multivariate technique Partial Least Squares Correlation (PLSC) (McIntosh, et al., 1996; Wold, 1982) to estimate possible covariance with GM properties and the distinctive aspects of each white matter metric. To examine the grey matter-white matter covariance patterns, six separate PLSC analyses were conducted: one for GM thickness and its relation to WM FA, one for GM thickness and its relation to WM MD, and the same for surface area and FA and MD, and volume and FA and MD. Because volume is composed of both thickness and surface area, we chose to focus on each aspect of volume in this report, with volume-FA and volume-MD multivariate results presented as Supplementary Information. Separate data matrices were created; one for each diffusion metric (FA and MD), as well as one for each GM property (grey matter thickness, surface area or volume). By submitting one data matrix composed of white matter values, extracted from the standardized diffusion images, and a second data matrix of grey matter thickness, surface area or volume, extracted from the Freesurfer cortical parcellations, covariance between the two structural imaging techniques emerge. The table composed of white matter metrics was created by registering each individual’s diffusion images to a standardized white matter space, followed by reorganizing of the 3D matrix into a vectorized whole brain. Once combined across participants, the data matrix consisted of 186 participants organized in rows with vectorized whole brain voxels in columns. This allowed each column within the data matrix to represent the same point in the brain for each participant. The table composed of grey matter thickness (or surface area or volume) consisted of the same 186 participants organized in rows with grey matter thickness (or surface area or volume) for each ROI organized along the columns. Additionally, the potential effects of sex were residualized from each data table before analysis.

Once raw data values were organized in their respective tables, columns were centered and normalized such that the sum of squares within a column was equal to one. A correlation matrix was computed from the normalized columns of the data tables and orthogonal factors were decomposed using singular value decomposition. This resulted in a matrix of left singular vectors (or saliences) for the white matter data, a matrix of right singular vectors for the grey matter data, and a diagonal matrix of singular values (Abdi and Williams, 2013; Krishnan, et al., 2011). The original observations were then projected onto their respective saliences to observe their contribution to the derived components. These linear combinations of the original variables form the latent variables, which express the covariance between the two data tables (Krishnan, et al., 2011).

To test for statistical significance of the derived components, permutation resampling was used to create new, rearranged matrices from our original data that were then submitted to PLSC. Specifically, 10,000 permutations were run to reorganize the participants (data rows) while holding the brain variables (data columns) constant. Submitting the permuted data to PLSC results in a distribution of new singular values which are used as a null hypothesis to test against the original singular values (Krishnan, et al., 2011; McIntosh and Lobaugh, 2004). To test the reliability of the data elements (grey matter ROIs and white matter voxels) an inferential bootstrap technique was used to create entirely new sets of data by sampling with replacement. Through bootstrap resampling, bootstrap ratios (BSR) were derived by dividing the mean of the distribution by its standard error. The larger the BSR the more it is significantly stable (Abdi and Williams, 2013; Ferreira, et al., 2015; McIntosh and Lobaugh, 2004). Bootstrap ratios function similar to *z*-scores and are considered significant when the ratio is greater than 1.96, which equates to a *p*-value of .05. Due to the large number of voxelwise comparisons computed in this dataset, we selected a more conservative BSR threshold of ±3, which equates to a *p*-value of .0027, as a cutoff to interpret stability of the observed saliences. Because many of our findings survived the inferential analysis, interpretations rely instead on the magnitude of the BSR relative to other data points. The PLSC analysis was run using TExPosition, part of the ExPosition analysis package (Beaton, Fatt, & Abdi, 2014) within RStudio, while the inferential analysis was run using in-house code (MATLAB 2012b, The MathWorks Inc., Natick, MA) based on the Welford/Knuth algorithm (Welford, 1962).

## Results

### Univariate

Univariate analyses demonstrated that much of the grey and white matter tissue is susceptible to the effects of aging, beyond the effects of sex. Grey matter measure age results are summarized by region in Figure 1 and white matter diffusion metric age results are illustrated by voxelwise plots in Figure 2.

#### Grey Matter Measures

Standardized parameter estimates for age effects on volume, cortical thickness, and surface area are plotted together by region in Figure 1. Each model included both a quadratic and linear age term to test for a non-linear effect of age. In cases where the non-linear term was not significant it was removed and only the linear effects were modeled. There was a significant, negative, linear effect of increasing age on volume in all grey matter regions measured (illustrated by the first bar in each ROI in Figure 1). For three of these regions (posterior cingulate, temporal pole, and transverse temporal) the age effects were significantly better modeled with a quadratic fit. For all three GM metrics a significant quadratic fit is indicated with an asterisk in Figure 1. Similarly, increasing age was associated with significantly thinner cortex in all grey matter regions, with the exception of the caudal anterior cingulate gyrus (illustrated by the middle bars in Figure 1). This relationship was linear in most regions, but quadratic in the caudal anterior cingulate, entorhinal, inferior temporal, lateral occipital, medial orbitofrontal, superior parietal, and temporal pole. In contrast, surface area measurements were found to be significantly smaller with increasing age in 28 of the ROIs, but not for the other six (insula, paracentral, entorhinal, rostral anterior cingulate, medial orbitofrontal, and superior temporal), suggesting that regional surface area is more resilient to the effects of aging (see last bar in each ROI in Figure 1). Only the lingual gyrus and transverse temporal regions fit a quadratic age trajectory better than a linear. Comparing across the standardized parameter estimates also reveals regional trends among grey matter measures. The strongest linear effects of age on both cortical thickness and volume were found in frontal lobe regions, most notably the parstriangularis, parsopercularis, and superior frontal gyrus, followed closely by parietal regions: precuneus, supramarginal gyrus, and angular gyrus, and some temporal regions (superior temporal, middle temporal and fusiform gyri). Very few occipital and cingulate regions showed strong age effects for volume or thickness measures. In contrast, when comparing surface area measures, the fusiform, posterior cingulate, middle temporal, banks of the superior temporal sulcus, and the parahippocampal gyrus were all among the most highly negatively associated with age.

**Figure 1.**
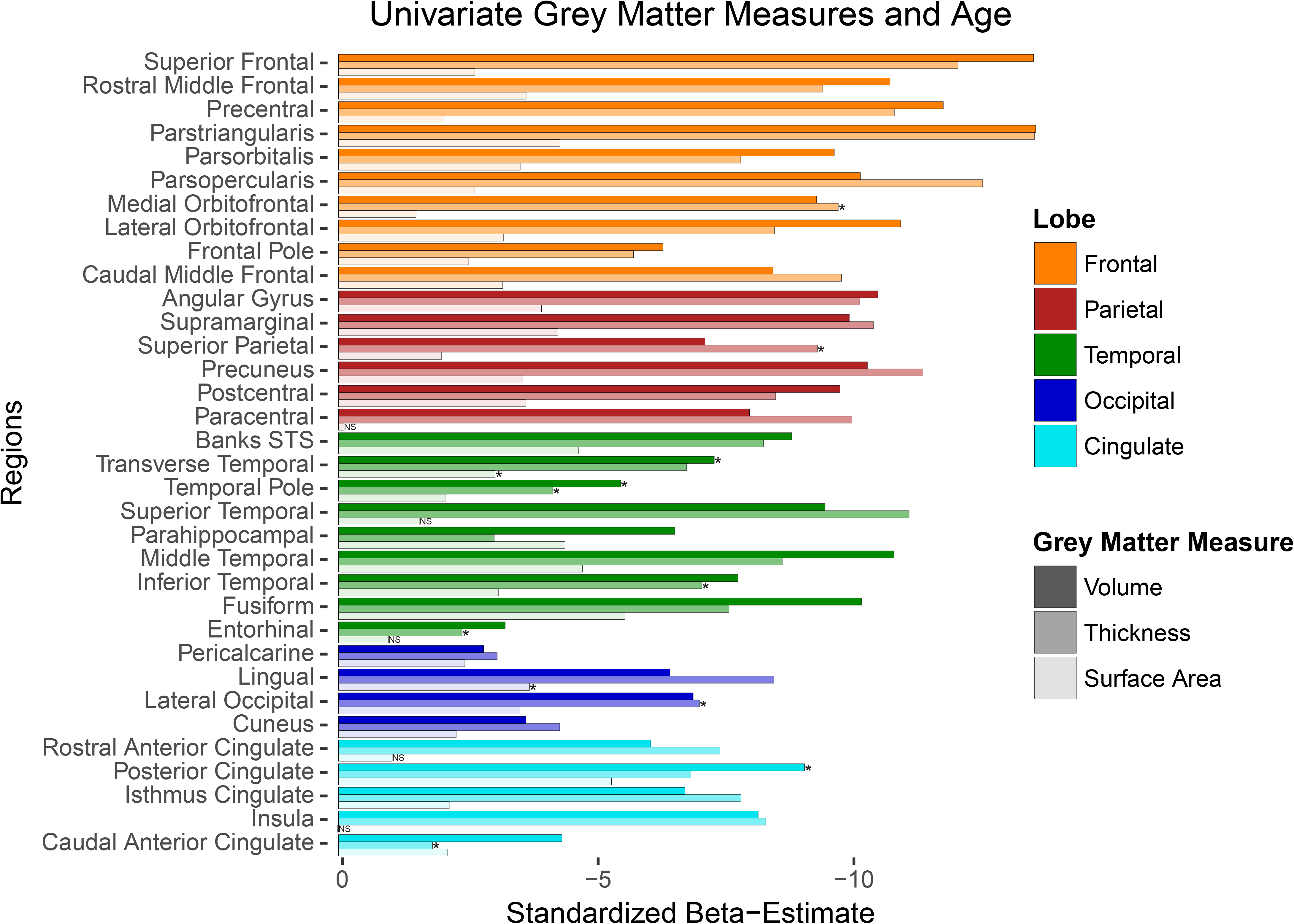
Univariate results for age differences in grey matter measures of volume, thickness, and surface area by region. Values represent standardized beta-weights from regression models containing age and sex as predictors. Larger estimates represent greater negative effect of age. Note. STS = Superior Temporal Sulcus. Bars with asterisks indicate measures that show significant quadratic effects of age (beyond linear). Bars with NS indicate measures with neither linear nor quadratic age effects.

#### White Matter Measures

Both white matter FA and MD measures indicated extensive age-related decreases in white matter health throughout the brain (see Figure 2). Over 86.2% of WM voxels analyzed evidenced a negative association between age and FA, and over 80.5% showed a positive association between age and MD. Only the center and occipital projections of the splenium of the corpus callosum and the inferior-most projection fibers extending into the cerebellum and spinal cord from the brainstem showed no relationship with age in either white matter metric (see green voxels in Figure 2). Unique effects of age on FA (i.e., not found on MD) were seen in the cerebellum, hippocampal projection of the cingulum, posterior cingulum bundle, projections into the superior-most parts of the frontal and parietal gyri from both the U-fibers of the corpus callosum and the corticospinal tract, and the posterior-most projections of the inferior longitudinal fasciculus, inferior-frontal occipital fasciculus, optic radiations, and splenium (see Figure 2, panel A). These unique effects with FA were mixed between linear (yellow voxels) and quadratic (blue voxels for quadratic, and pink voxels for linear/quadratic overlap) effects with no obvious regional specificity to the shape differences noted. Unique effects of age on MD (i.e., not seen for FA) were observed in medial portions (viewed sagittally) of the corona radiata and corticospinal tract, and the cingulum bundle body (see Figure 2, panel B), most of which demonstrated a quadratic association with age (illustrated in blue/pink voxels). The remaining portions of the cerebral white matter exhibited significant age-related association with both FA and MD measures including the superior longitudinal fasciculus, superior frontal occipital fasciculus, inferior longitudinal fasciculus, large anterior and posterior portions of the inferior frontal occipital fasciculus, remaining portions of the corpus callosum, uncinate fasciculus, and fornix (see Figure 2, panels A and B). Most of these regions demonstrated a quadratic relationship with age (blue/pink voxels), especially between MD and age (panel 2B), while FA showed a linear relationship in medial parts of the corpus callosum, the center and anterior projections of the genu, and in some of the U-fibers projecting to the frontal lobe gyri (yellow voxels in panel 2A).

**Figure 2.**
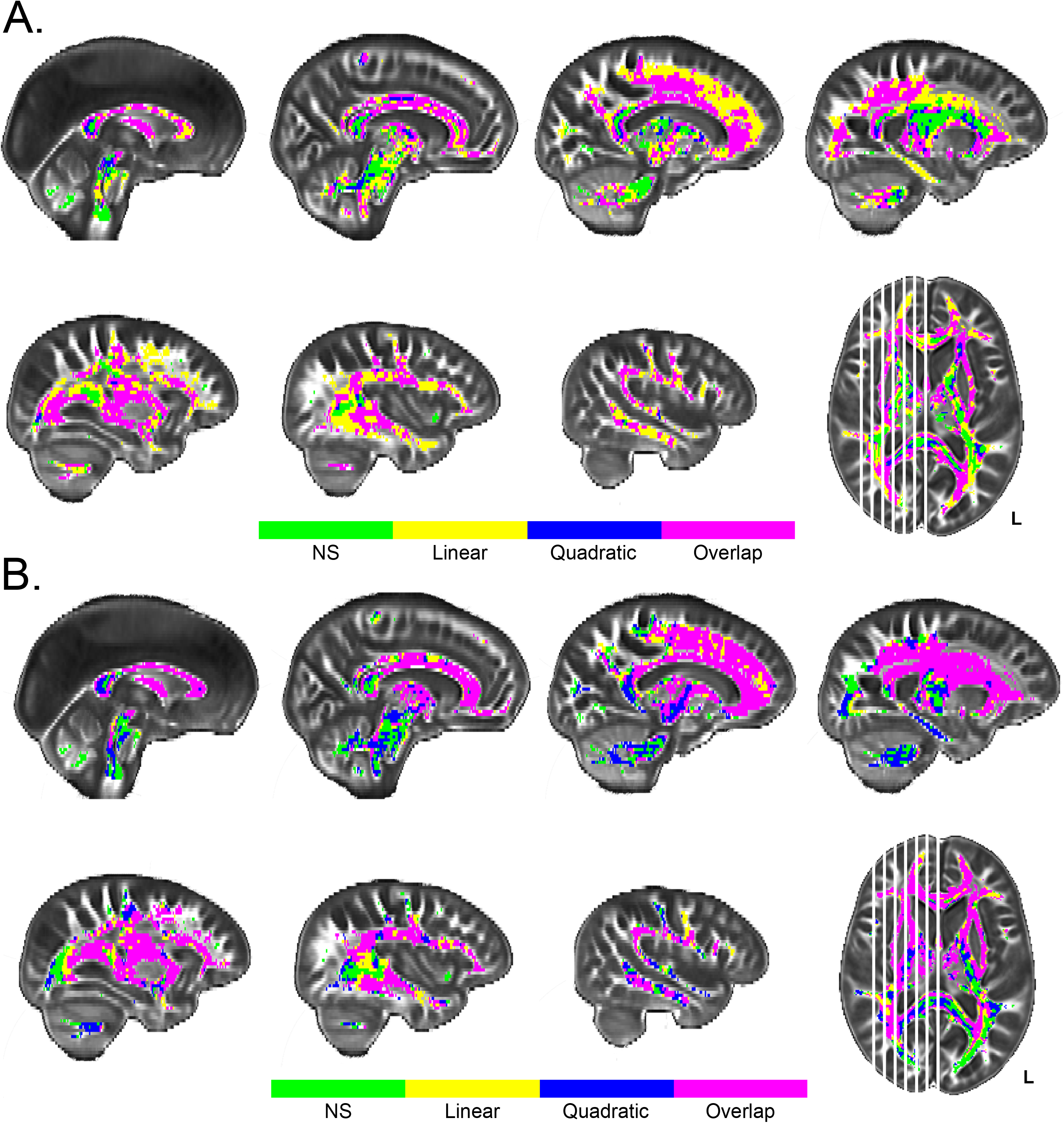
Univariate results for age differences in white matter FA (A) and MD (B). Panel A depicts regions with significant effects of age projected onto sagittal slices. Linear effects of age on fractional anisotropy (FA) are depicted in yellow, quadratic effects of age in blue, and their overlap in pink. Panel B depicts regions with significant age effects on mean diffusivity (MD) following the same voxel color scale as for FA in panel A. Plotted voxels represent significant voxels from Threshold-Free Cluster Enhancement (TFCE) raw test statistics that survived permutation correction. In both panels, green voxels represent regions of age invariance.

Spatial Gradient Patterns. To systematically depict any gradient patterns of age effects on FA and MD, the strength of the association between age and each WM metric is plotted across the slices of the brain in Figure 3. Examination of the plots in Figure 3 suggests that traversing in either the posterior-to-anterior direction or the inferior-to-superior direction there is a strong age gradient effect demonstrated by the increasingly strong association between both age and FA (in Figure 3, panel A) and age and MD (in Figure 3, panel B) when moving anteriorly and moving superiorly across the brain. Interestingly, there is also a strong medial-to-lateral gradient in the relationship between age and MD that decreases when moving from mid-sagittal toward the lateral most slices, which is reversed with FA (Figure 3).

**Figure 3.**
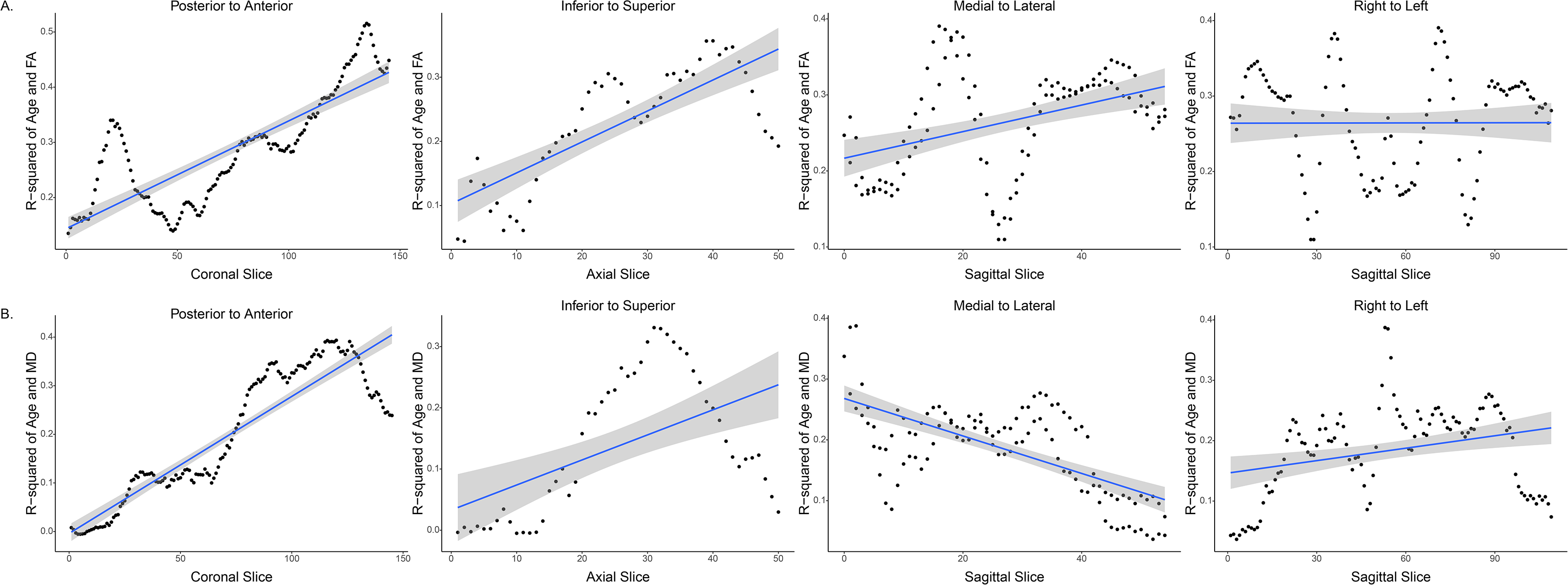
Strength of relationship between age and white matter metric plotted across brain slices illustrating spatial age gradients. Panel A plots fractional anisotropy-age correlations and Panel B mean diffusivity-age correlations across slices in four directions. There is a strong, posterior-anterior and inferior-superior gradient increase in the strength of the effect of age on both FA and MD. There is also a medial-to-lateral (sagittally from midsagittal to parasagittal across both hemispheres) gradient of age effect on MD and FA. Hemispheric (laterality) differences, sagitally from right to left, were minimal.

### Multivariate

Separate partial least squares correlation analyses were applied to examine how estimates of neuronal degradation were associated across the lifespan: two each (one for FA and one for MD) for each of the grey matter regional measures (cortical thickness, surface area, volume) for a total of 6 analyses. The univariate findings suggested that age-related variance in volume was driven primarily by thickness values, and that surface area was relatively age-invariant. Therefore, the presentation of multivariate results is primarily focused on cortical thickness with surface area and volume PLSC results presented as Supplemental Material.

### Cortical Thickness

#### Inferential analysis

The first set of two analyses combined GM thickness and WM FA (PLSC-FA), while the other combined GM thickness and WM MD (PLSC-MD). Each analysis revealed a single significant component that explained 94.42% and 91.48% of the variance in the data, respectively for PLSC-FA and PLSC-MD, at a *p*-value of *p* < 0.0001. No other components resolved, from either analysis, explained a significant amount of variance. An inferential battery was performed on the saliences of each component to establish reliability of these results. After averaging across each hemisphere, all grey matter regions, in both analyses, survived a bootstrap ratio cutoff of ±3 (*p* = .0027), with the exception of the entorhinal cortex in PLSC-FA, pericalcarine cortex in PLSC-MD, and caudal anterior cingulate and parahippocampal gyrus in both analyses (Figure 4A and Figure 4B).

**Figure 4.**
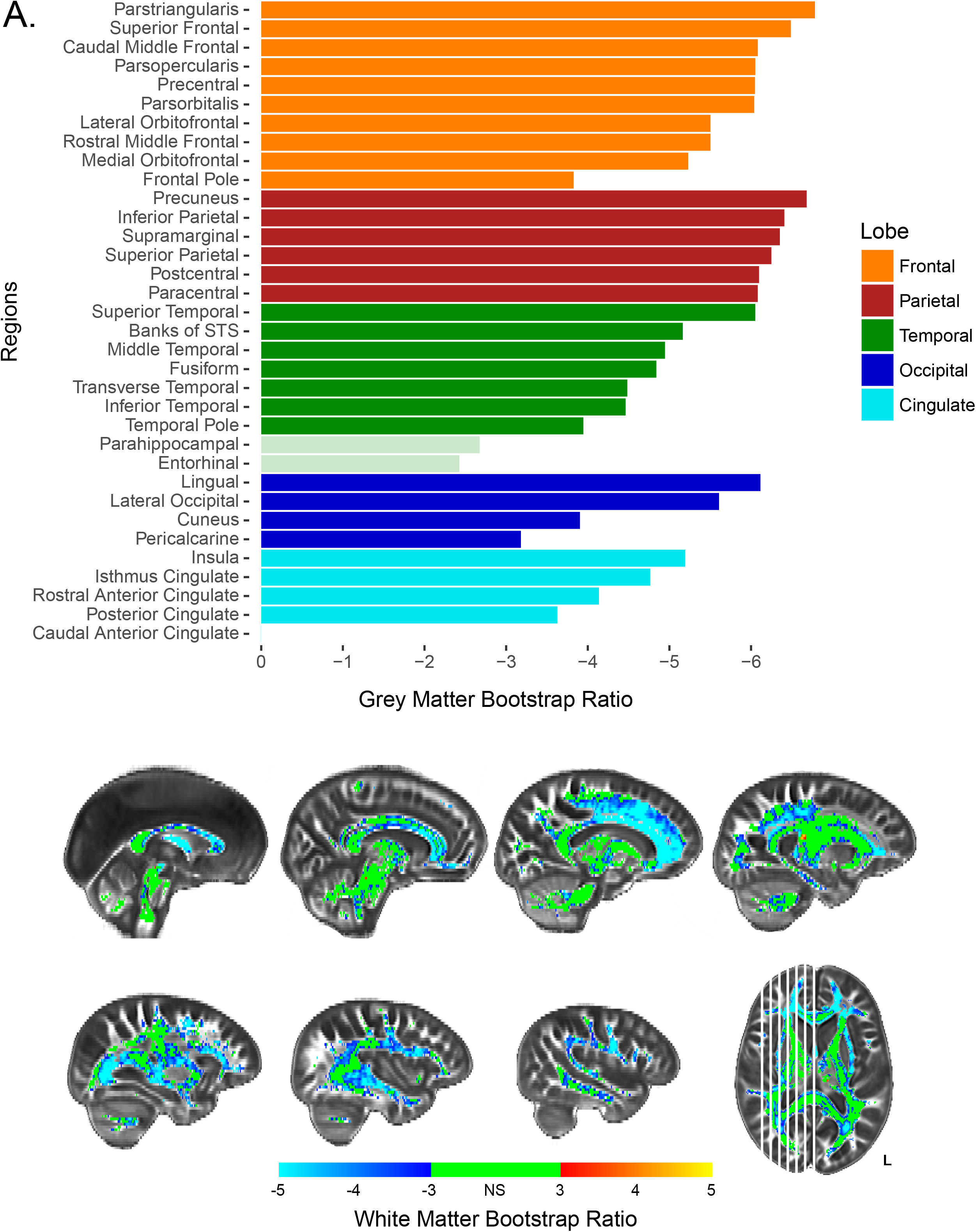
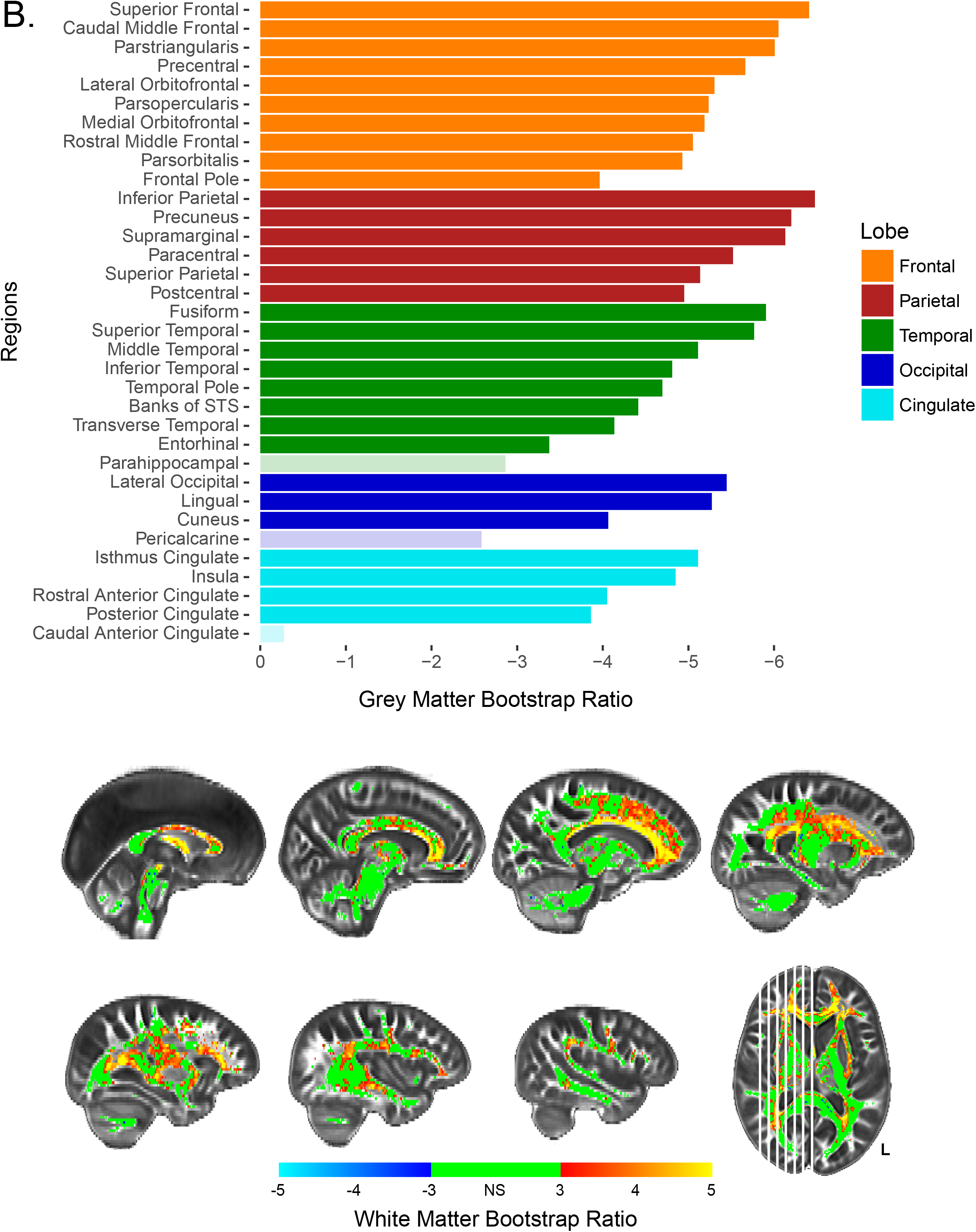
Bootstrap ratios for PLSC-FA (A) and PLSC-MD (B). Barplots show bootstrap ratios for GM regions, while brain images show bootstrap ratios for WM voxels. Bootstrap ratios below an absolute value of 3 (which corresponds to *p* < .0027) are considered non-significant and are faded in GM barplots, or green in WM voxelwise brains. Note. STS = Superior Temporal Sulcus.

In PLSC-FA, the most stable GM saliences were those of the parietal regions, which all obtained BSR > 6 (*p* < .0001). All frontal regions obtained BSR > 5 (*p* < .0001), with the lone exception being the frontal pole. Additionally, while most temporal, occipital, and cingulate regions were the least stable, the lingual, superior temporal, and lateral occipital gyri all obtained high bootstrap ratios demonstrating high importance in the analysis (Figure 4A). Similar to grey matter thickness, most FA voxels survived an inferential bootstrap ratio cutoff of 3, with the few exceptions being medial portions of the corticospinal tracts, projections from the brainstem, and the splenium of the corpus callosum (Figure 4A). The most significant FA voxels, obtaining a BSR ≥ 5, were white matter connections typically thought to traverse frontal and parietal regions. These include the projections from the genu and body of the corpus callosum including the U-fibers extending into frontal and parietal gyri, anterior portions of the SLF and superior frontal occipital fasciculus, and the fornix. In this analysis, both GM and WM exhibit negative BSRs, which indicates a positive association between the two tissues—as thickness in frontal and parietal cortex decreased, FA in largely frontal and parietal white matter decreased.

In PLSC-MD, the most stable GM saliences were both the frontal and parietal regions as all regions obtained a BSR around 5 or above (p < .0001), with the exception of the frontal pole. Similarly, most temporal, occipital, and cingulate regions were among the least stable. However, the fusiform, superior temporal, lateral occipital, and lingual gyri were each important contributors to the analysis (Figure 4B). Results with WM MD were similar to those with WM FA. The most stable white matter voxels were those of both the frontal and parietal regions, although most frontal and parietal regions showed less stability with MD than they did with FA. Interestingly, many temporal and cingulate regions were shown to be more stable with MD than with FA. The most significant MD voxels, obtaining a BSR > 5, (p < .0001), were the centers of the genu and body of the corpus callosum as opposed to the projections and U-fibers observed with FA (Figure 4B). Additionally, the medial portions of the corona radiata (superior to the thalamus but inferior to the cortical gyri), the anterior thalamic radiations, and the fornix, also were shown to share the most variance with grey matter thickness. Opposite signs of the GM and WM BSRs in this analysis indicates a negative relationship between the two measures—as grey matter thickness decreased in frontal and parietal cortex, MD increased in medial white matter regions.

#### Latent analysis

PLSC provides “brain scores” for each participant which describe the degree to which an individual expresses the white matter and grey matter latent variables from the model (Krishnan et al., 2011). A brain score near zero indicates that the variance attributed to the model from a given data point contributed less informative or influential information to resolving the component, or was highly similar to the mean participant for that measurement. In contrast, the greater the magnitude of the brain score the more influential a given data point was in the overall co-variance structure. Figure 5, panel A illustrates how the observations (each participant in the sample) contributed to GM thickness (y-axis) and WM FA (x-axis) of component 1 while Figure 5, panel D shows these same observations and the contributions of GM thickness (y-axis) and WM MD (x-axis) of component 1. Observations are colored by age group, for visualization purposes, to illustrate how component 1 can be interpreted in the context of an age gradient.

**Figure 5.**
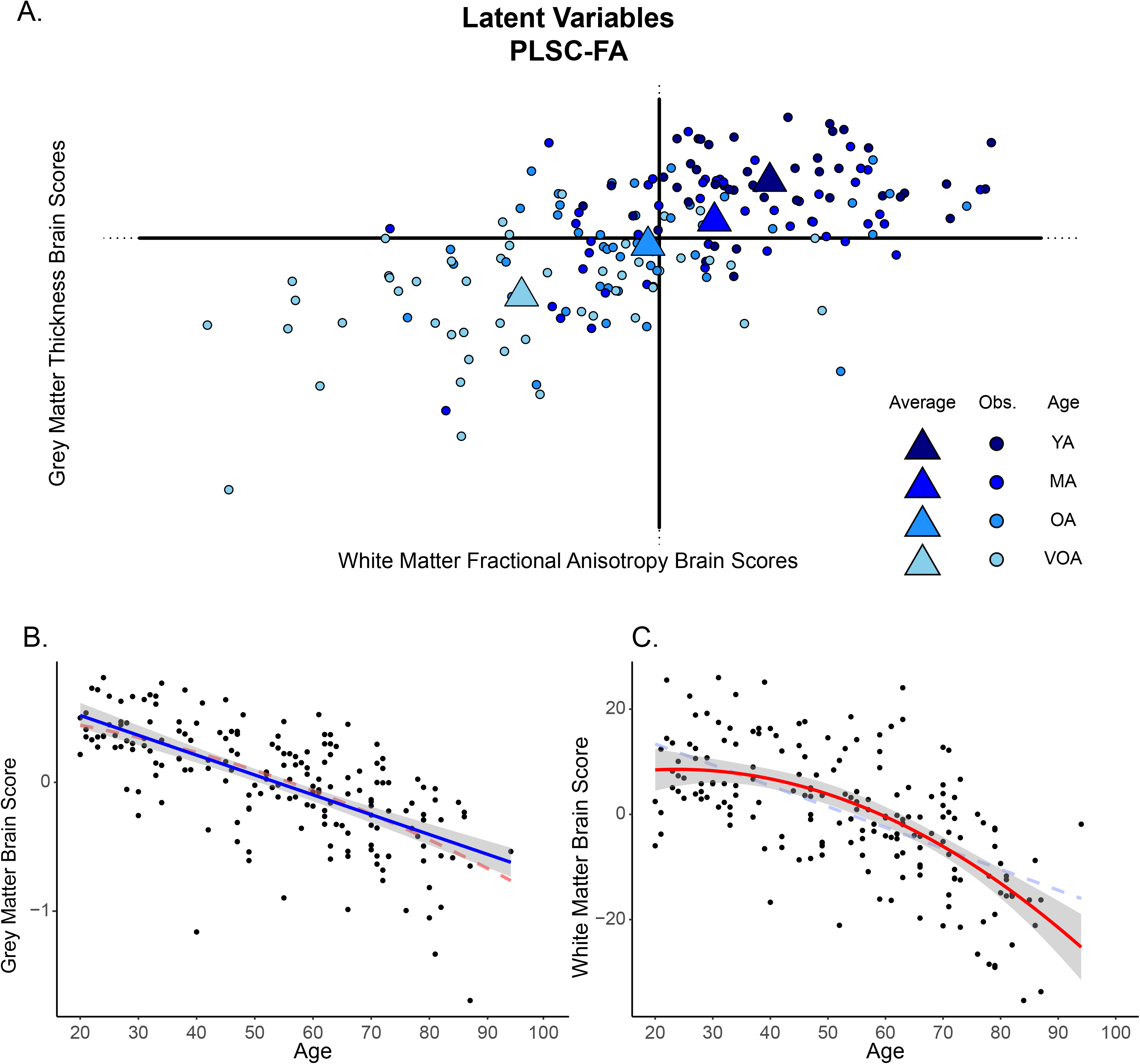
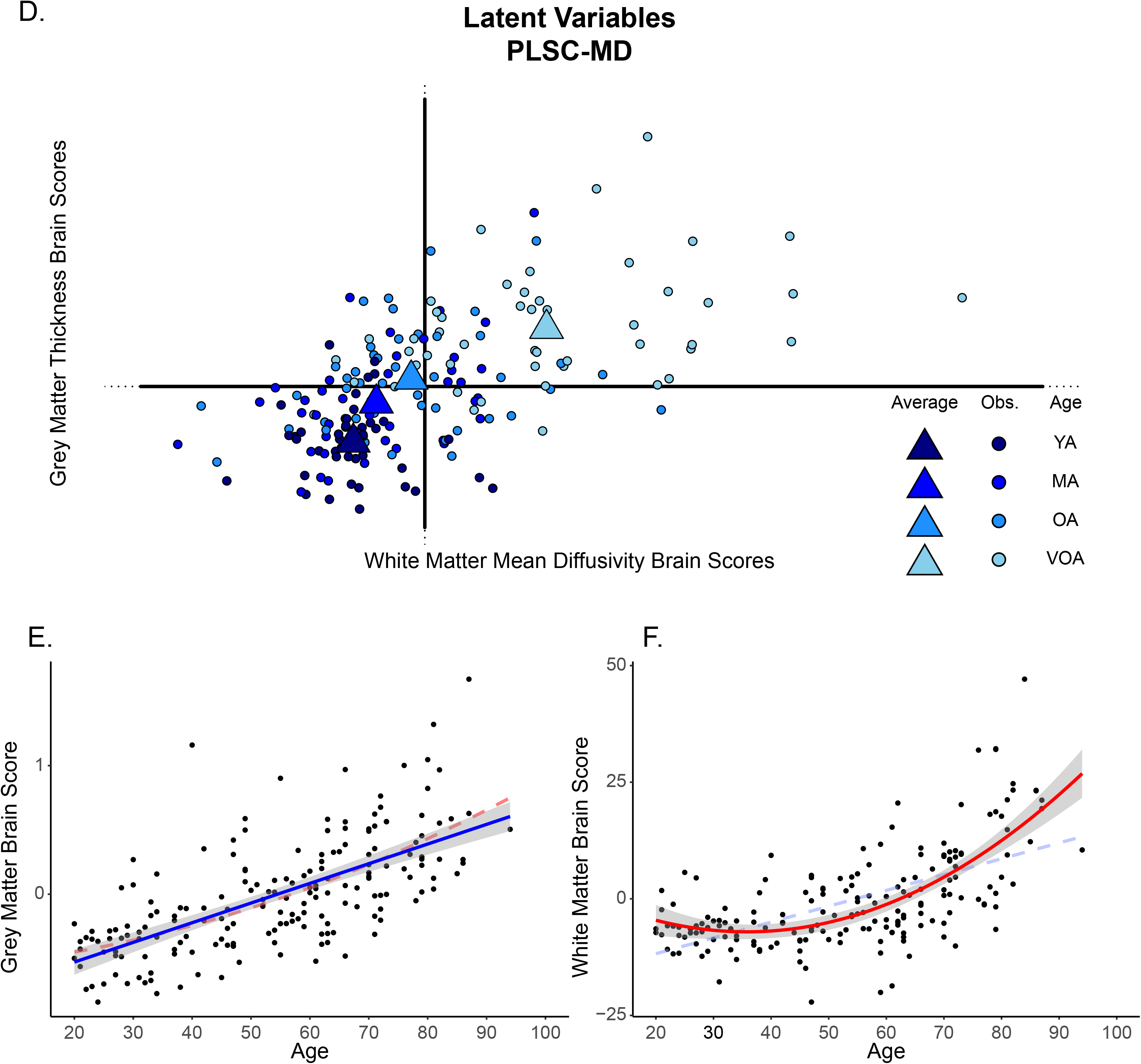
Latent variable projections and the relationship of each brain score with age for (A-C) PLSC-FA and (D-F) PLSC-MD. For the PLSC-FA analysis, grey matter brain scores decrease linearly with age demonstrating that data from younger adults drive the positive side of component 1, while data from older adults drive the negative side of component 1 (illustrated in panel B). This association is also evident for white matter brain scores in a nonlinear fashion (i.e., decelerates with age; panel C). For the PLSC-MD analysis similar linear (for grey matter in panel E) and quadratic (for white matter in panel F) relationships with age are found. Scatter dots represent individual participants, triangles represent the mean of an age “group”, and blue color fade from dark to light blue represents younger to older adults, respectively to illustrate the age effect. Both linear and nonlinear regression lines are illustrated on the scatterplots, with the more significant fit denoted with bold solid lines, and the lower fitting line denoted by faded dashed lines. Note. YA= Younger Adults; MA = Middle-Aged Adults; OA = Older Adults; VOA = Very Old Adults.

Plotting the WM and GM brain scores together shows how the age of the participants describes the resolved component as there is a relatively linear progression from the youngest adults (darkest blue), to oldest adults (lightest blue), across the first component in each PLSC. Additionally, the brain score magnitude of both the young adults and very old adults (i.e., the distance from the origin), shows that they are driving the variance and contributing to the significance of the component. To test this relationship more directly, regression curves are depicted demonstrating the association between GM (Figure 5 panels B, E) and WM (Figure 5 panels C, F) brain scores from each PLSC and participant age. There is a significant linear effect of each brain score and age; PLSC-FA: WM ~Age (*F*(1,184) = 111.7, *p* < .001), GM ~ Age (*F*(1,184) = 143.9, *p* < .001); PLSC-MD: WM ~Age (*F*(1,184) = 106.4, *p* < .001), GM ~ Age (*F*(1,184) = 141.1, *p* < .001). Given the strength of these associations, the PLSC results can be interpreted relative to age by combining information from the two salience plots with the chronological information provided from the latent mapping of the brain scores. Interestingly, there is also a significant quadratic effect of age for each WM brain score; PLSC-FA: WM ~ quadratic Age (*F*(2,183) = 64.72, *p* <.001); PLSC-MD: WM ~ quadratic Age (*F*(2,183) = 81.14, *p* < .001), but not for GM brain scores.

The PLSC-FA saliences for both the grey matter ROIs and the voxel-wise white matter FA estimates are on the same side of the component structure (evidenced by the negative saliences leading to negative BSRs in Figure 4A), suggesting a positive relationship between grey matter thickness and white matter microstructure (e.g., as thickness increases, FA values increase). The plot of brain scores for this analysis reveals that older adults, plotted in the bottom left quadrant (i.e., negative brains scores for both grey and white matter latent variables), demonstrate a negative relationship with the coupled variance of grey matter thickness and white matter structural measures (Figure 5A). In contrast, the youngest adults fell to the upper-right quadrant of the first component demonstrating a positive relationship with the coupled variance of grey matter thickness and white matter structural measures. Together, these results indicate that the age gradient aligns with the coupled variance in structural measures such that the youngest adults exhibit the expected positive association between grey matter thickness and FA estimates, while older adults are better characterized by coupled decreases in structural integrity.

In PLSC-MD, a similar pattern emerges among the coupling of structural integrity measures and age. The saliences for the grey matter ROIs are along the left side of the component structure (leading to negative BSRs in Figure 4B), while the saliences for the voxel-wise white matter MD are along the right (leading to positive BSRs in Figure 4B), suggesting a positive relationship between grey matter thickness and white matter microstructure (Figure 5, panel D). This is due to the fact that higher MD is typically associated with poorer structure in healthy adults, thus a sort of “double-negative” association emerges between the two structural measures. When examining WM and GM brain scores, the oldest adults are projected to the upper right quadrant of the first component which is characterized by positive GM brain scores (opposite the negative GM saliences) and positive WM brain scores (same as the positive WM saliences) (Figure 5D). This demonstrates a negative relationship with the coupled variance in structural decline of grey matter thickness and white matter structural measures. Taken together, these results also indicate that the age gradient aligns with the coupled variance in structural measures (i.e., the youngest adults exhibit a positive relationship, while older adults exhibit a negative relationship).

Anatomical specificity is evident due to the magnitude of the saliences and latent projections, or the distance each data point is from the origin. In the PLSC-FA analysis, grey matter thickness measures in the parietal and frontal regions, specifically the superior frontal, parstriangularis, precuneus, angular gyrus, and supramarginal gyri are driving the observed variance (Figure 4A). In other words, the largest covariance between thickness and white matter FA, or greatest coupling, appears in these association cortices. Similarly, the white matter saliences for FA, which show the greatest coupling with grey matter thickness, are the voxels in the brightest blue (Figure 4A). These consist of a large amount of frontal voxels such as the genu and body of the corpus callosum, anterior portions of the cingulum bundle, U-fibers extending from anterior portions of the corpus callosum to frontal and parietal gyri, the superior longitudinal fasciculus, and the fornix. In contrast, thickness values from the parahippocampal and entorhinal regions in the temporal lobe share little variance with FA values, while the caudal anterior cingulate shares no variance with FA values. White matter FA of the more posterior and inferior regions, including the splenium of the corpus callosum, projections from the corticospinal and corona radiata, anterior thalamic radiation, projections into the occipital lobe and the brainstem, and cerebellar fibers share very little variance with measures of thickness.

The PLSC-MD analysis reveals that grey matter thickness measures in the parietal and frontal regions, specifically the superior frontal, caudal middle frontal, angular gyrus, precuneus, and supramarginal, are driving the observed covariance (see Figure 4B). Similarly, the white matter saliences for MD, which show the greatest coupling with grey matter thickness, are composed of a large number of frontal voxels such as the genu and body of the corpus callosum, U-fibers extending from anterior portions of the corpus callosum to frontal gyri, medial portions of the corona radiata and corticospinal tract, and the fornix (Figure 4B). In contrast, thickness values from the parahippocampal, pericalcarine, and caudal anterior cingulate, share little variance with MD values. White matter voxels of the more posterior and inferior regions, including the splenium of the corpus callosum, the inferior longitudinal fasciculus, projections into the occipital lobe and the brainstem, and the cerebellum, share little variance with measures of thickness.

### Surface Area

#### Inferential Statistics

PLSC models were also conducted for regional surface area and FA and MD. Surface area demonstrated fewer and less robust covariance with white matter, as surface area is less susceptible to the effects of aging, compared to cortical thickness. Surface area PLSC results are provided as Supplemental Information. Supplemental Figure S1, panel A illustrates that there is significant covariance with FA and surface area in rostral and caudal middle frontal and superior frontal gyri, in all parietal regions but paracentral, in middle temporal, banks of the superior temporal, and fusiform gyri, in all occipital regions, and in only the posterior portion of the cingulate gyrus.

Supplemental Figure S1A also illustrates the white matter voxels where FA covaries with surface area. These voxels are sporadic, but tend to cluster in the projections of the corpus callosum, including the genu and u-fibers extending from the body of the corpus callosum to frontal and parietal gyri, as well as the frontal projections of the IFOF, the cingulum body, the SLF, and fornix.

For the covariance between surface area and WM MD, Supplemental Figure S1 panel B illustrates significance in the caudal and rostral middle frontal and superior frontal gyri, in postcentral, supramarginal, and precuneus, in fusiform, middle temporal, transverse temporal and banks of the superior temporal gyri, in all of occipital regions, and in the posterior portion of the cingulate gyrus. Supplemental Figure S1B also illustrates the white matter MD voxels where there is significant covariance with surface area. These voxels are even more sporadic than in the surface area analysis with FA, including the genu and some projections of the corpus callosum to the frontal lobes, as well as the fornix and anterior IFOF.

#### Latent Analyses

Supplemental Figure S2A provides “brain score” plots for the covariance between surface area and WM FA in terms of latent components. As in the cortical thickness results shown in Figure 5, the first component is interpreted as age. Supplemental Figure S2 B/C illustrates the strength of the brain score for grey matter and white matter, respectively, and age. Supplemental Figure S2 D, E, and F displays the same information for WM MD.

### Grey Matter Volume

#### Inferential Statistics

We also include as supplemental information the PLSC results for the covariance between grey matter volume and WM FA and MD. Because volume is the product of surface area and cortical thickness, it stands to reason that its covariance would be similar to one of its driving components. Indeed, the covariance patterns for volume appear similar to the cortical thickness covariance and distinct from the surface area pattern. Supplemental Figure S3 provides regional volume BSR plots and WM maps for FA (Supplemental Figure S3A) and for MD (Supplemental Figure S3B). Significant covariance between GM volume and WM FA was observed in every region except the caudal portion of the anterior cingulate gyrus. GM volume and WM MD covariance was also significant for every region except caudal anterior cingulate. In addition, significant WM voxels mirrored the regional distribution in the thickness results with large areas of frontal and parietal WM dominating the covariance for both FA and MD.

#### Latent Analyses

“Brain score” plots for the covariance between cortical volume and WM FA in terms of the first latent component, with age coded across the plot are shown in Supplemental Figure S4A. The relationships between each brain score and age are shown in Supplemental Figure S4 B and C, while the results for volume and WM MD covariance are provided in Supplemental Figure S4 D, E, and F.

## Discussion

Neuronal degradation in the aging brain is a complex biological process involving changes in both grey and white matter tissue. MRI metrics, serving as a proxy for these tissue compartments, show differential effects of normal aging and are generally investigated separately. Here, we report traditional univariate findings of normal aging on grey matter thickness, surface area, and volume as well as fractional anisotropy and mean diffusivity differences in white matter across the adult lifespan. We replicate previous findings to demonstrate how each tissue type is associated with age. We then report results using a multivariate technique to investigate the effects of aging on both tissue types simultaneously to demonstrate how regional tissue degradation has both shared and unique aspects across the lifespan, elucidating this relationship in a way univariate methods cannot. We found that variance in the data follows an aging trajectory evidenced by intact structure in early adulthood, followed by differential degradation, in regards to both tissue type and anatomy, throughout mid-to late-adulthood.

### Univariate Findings

Previous research illustrated that age-related declines in grey matter follow a “patchwork” pattern of regional specificity that roughly follows a “last-in first-out” gradient of vulnerability such that structures that evolve and/or mature earliest are the most resilient to the processes of aging, and likewise, those with the most protracted development are the most vulnerable to the aging process (Raz, 2000; Salat, et al., 2004). Although we did not explicitly test this hypothesis by directly comparing early and late developing regions, the univariate results in the current study largely replicate and expand on this idea. The cortical thickness and volume measures with the highest aging parameter estimates are all located in the frontal lobe. Other high parameter estimates were those of the parietal regions, while the only temporal regions with high estimates were that of the superior and middle temporal gyri, which are known for contributions to higher order cognitive processing and demonstrate susceptibility to aging (Fjell, et al., 2009; Van Petten, et al., 2004). A regional gradient is evident such that the frontal regions have higher parameter estimates than the majority of the occipital, temporal, and cingulate regions, meaning that these association cortices explain significantly more age-related variance. Interestingly, surface area measures with the highest age-related parameter estimates include occipital and temporal regions, some of which are reportedly more highly associated with age than with thickness or volume (Dickerson, et al., 2009). However, age-related parameter estimates of surface area are modest relative to volume and thickness, suggesting that while surface area does show differences with age, it is not nearly as age-vulnerable as other measurements (Lemaitre, et al., 2012). As suggested by Lemaitre et al., it might be the case that surface area measurements are more sensitive to the loss of intracortical white matter.

Alternately, cortical thickness has been suggested to reflect, during development, dendritic arborization and pruning in grey matter (Huttenlocher, 1990), and myelination differences at the interface of grey and white matter tissue compartments (Sowell, et al., 2004). During development surface area is related to the maintenance and division of progenitor cells (Chenn and Walsh, 2002). However, both cortical thickness and surface area development are related to neuronal migration and minicolumn organization (Rakic, 1988). Surface area may reflect the density of minicolumns in cortex (Rakic, 1988), whereas cortical thickness reflects the size and density of cells, including those that make up minicolumns (Rakic, 1988), at least in animal models of development. Cortical thickness appears to be most impacted developmentally by alterations in myelin, reduction in size or number of neurons, and changes to the synaptic environment (Sowell, et al., 2003; Sowell, et al., 2004), which are the hallmarks of age-related decline, suggesting a biological basis for a retrogenesis-like recapitulation of development in aging.

In univariate analyses, measures of white matter FA and MD also show strong associations with age. Analyses revealed linear gradients of age-differences across much of the myelinated area, indicating that regional specificity might be more complex than a simple anterior-to-posterior gradient or last-in first-out framework. While anterior regions did evidence strong age-related associations for both FA and MD compared to posterior regions (lower FA and higher MD while moving toward anterior voxels), there were also strong medial-to-lateral effects such that MD evidenced weaker age associations while moving from mid-sagittal to more lateral slices. Age-related associations with FA appear to reflect an overall disruption in WM microstructure at the tract centers, which remained stable in large white matter bundles such as the corpus callosum, yet declined in smaller offshoots that approach lateral gyri. This finding corresponds with the idea that more primitive brain regions, which typically myelinate earliest in development and are more centrally located in medial portions of the brain, are typically more preserved in aging; as opposed to later myelinating association fibers, which connect cognitive centers across the superior and lateral surfaces, and exhibit greater age-related structural loss (Salat, et al., 2005; Ziegler, et al., 2010). Indeed, the latest myelinating regions are superiorly positioned – the supramarginal, superior frontal, and superior parietal – followed by inferior temporal, middle temporal, and superior temporal gyri, and the precuneus (Deoni, et al., 2015; Leipsic, 1901). In comparison, middle-development is characterized by myelination of the corpus callosum, and earliest to develop are the projection fibers (Brickman, et al., 2012; Stricker, et al., 2009). On the other hand, decreasing MD appears to be more specific to the edges of each tract, possibly representative of myelin degradation with age as larger WM bundles become less dense along the periphery. Demyelination and changes to myelin density might have an exacerbated effect on larger fiber bundles. Therefore, MD might be less sensitive to regional phylogeny or ontogeny and more sensitive to the size and/or density of fiber bundles. More research is needed regarding myelin density and how it is affected within individual tracts across developmental and aging processes.

### Multivariate Findings and Comparison to Univariate Results Pattern

Because the different processes of the neuron (cell body, axons, dendrites) likely age in conjunction, we sought to use multivariate analyses to expand upon current and previous univariate aging brain findings with a more biologically-relevant guiding model which simultaneously takes data from proxies of both the neuronal axon and cell body into account to gauge, at least with cross-sectional estimates, how these brain compartments may age together. Specifically, we utilized a partial least squares statistical technique to demonstrate how variance among grey and white matter estimates of neuronal integrity co-vary across the brain; and how this covariance varies across the lifespan. Comparisons among the two types of analyses show that multivariate techniques emphasize a coupled relationship of grey and white matter decline which is not evident from univariate analyses alone. While univariate analyses show the strongest effects of age and thickness to be in the frontal cortices, the co-variation with white matter FA reveals that the greatest coupled association is actually in frontal and parietal cortices and the frontal white matter connections. Additionally, occipital regions, which in the univariate analyses were among the least associated with age, are more highly associated with white matter metrics and aging once both tissues are considered together. In particular, aging of the lingual and lateral occipital cortices thickness show a strong coupling with both FA and MD in the multivariate analyses. In contrast, cingulate regions, which were only weakly affected by age in the univariate analysis, showed lesser association with both white matter FA and MD when analyzed together. Frontal regions, which were the most highly associated with age in the univariate analyses, still demonstrate high coupling with WM and age, albeit less than many parietal regions. These findings reveal that considering structural brain aging, as a whole, and the coupling among grey and white matter tissue, leads to different conclusions than an analysis which considers the neuronal processes as two unrelated tissue types. Previous studies suggest that correlations between cortical thickness across brain regions may be related to the function of the networks the regions participate in and to the underlying white matter connectivity (He, et al., 2007; Lerch, et al., 2006). This type of connectivity could then serve as a plausible explanation of how aging of one component of one region might affect downstream regions; however, this speculation requires longitudinal design to fully test.

### Regional Timecourse Specificity: Grey and White Matter Coupling

The current study not only demonstrated a coupled association of tissue integrity in a lifespan sample of healthy aging adults, but it also suggests a differential timecourse of aging within each tissue type, in accord with other cross-sectional studies (Fjell, et al., 2013; Raz and Rodrigue, 2006; Ziegler, et al., 2010). As discussed above, the relationship between structural measures is described by age such that the projection of the latent variable contributions map onto an age trajectory. However, despite this strong coupling of structural measures, regional aspects of grey matter cortices and white matter voxels differentially contributed to the age effects. Thus, it appears that different tissues types, despite their connectedness within a neuron, degrade at different points in the lifespan. This distinction is most notable in the relationship between frontal and parietal structural measures. In comparisons with white matter FA, parietal grey matter thickness measures were at the extreme end of the component. Therefore, the age-related thickness of the parietal cortices is not only sharing the most variance with estimates of white matter FA, but is also driven by the oldest adults. Frontal cortices also contributed heavily to this relationship and the component structure as a whole, but shares less variance with white matter FA than parietal regions. Interestingly, in analyses with white matter MD, this relationship is similar such that, as a whole, white matter MD shares the most variance with thickness measures of frontal cortices, followed by thickness measures from parietal regions. Therefore, there is an overall coupling of aging of grey matter thickness in the frontal and parietal regions with estimates of white matter microstructure, although regional specificity may be evidenced depending on which white matter metric is evaluated. Temporal and occipital regional thickness were even less representative in analyses with either white matter FA or MD, reflecting less coupling with age, with the exception of lingual and superior temporal cortices.

Our results indicate that similar regionally-specific features of white matter voxels emerge in both analyses. Frontal white matter voxels shared the most variance with age-related grey matter thickness in both analyses, specifically, white matter voxels spanning between frontal and parietal cortices such as the genu and body of the corpus callosum, the projections from the corpus callosum, and the anterior portions of the cingulum bundle. Comparisons between the two WM metrics revealed that FA demonstrated greater covariance (via higher bootstrap ratios) with GM thickness in some of the largest association fiber bundles, such as the anterior projections of the corpus callosum into frontal and parietal lobes, the frontal-parietal portion of the SLF, the cingulum bundle, and posterior projections of the splenium, ILF, and IFOF, compared to MD. Reshaping of the diffusion ellipsoid within these voxels, due to the loss of more thinly myelinated axons, without compromising the overall magnitude of water flow is plausible in these association fibers. In contrast, MD evidenced stronger age-related coupling with GM thickness in medial portions of the brain such as the corona radiata, cortical spinal tract, and anterior thalamic projection fibers. This could be the result of thickly myelinated, medially located bands of white matter, allowing for more water flow (due to less displacement) with aging, yet not necessarily reshaping the directionality of this flow (as would be measured by FA). This anatomical dissociation of tissue compartments is important with regard to age-related tissue alterations, as it suggests that certain tissue types are more susceptible to the aging process than others (Bartzokis, et al., 2012; Bartzokis, et al., 2004), and could yield insight as to how regionally differential neurons degrade according to different trajectories. Although imaging data with finer resolution than currently available at the cellular or neuronal level using longitudinal change data would be necessary to validate these ideas, one possibility based on the current results is that parietal neurons degrade from cell body to axons (e.g., as in Wallerian degeneration) whereas frontal neurons degrade from cell axons to body (e.g., transneuronal atrophy).

While the results from cross-sectional studies are limited in their ability to elucidate age progression within individuals, it is interesting to note that our findings are consistent with the general patterns of longitudinal studies (Rast, et al., 2017; Raz, et al., 2005). We found that grey matter regions that contribute the most variance, and display a negative trajectory with age, are parietal and frontal regions, while those regions contributing the least to this relationship with age are occipital and cingulate regions. This framework in which heteromodal, association cortices (forming higher-level cognition centers) evidence early decline and are associated with deteriorating cognition, with a relative preservation of primary sensory areas has been a proposed mechanism underlying cognitive decline patterns in healthy aging (Raz, 2000). Similarly, the strongest relationships in white matter degradation were evidenced by the negative age trajectory with areas throughout the anterior and superior white matter, typifying anatomical association connections between higher-order cognitive centers of frontal and parietal grey matter. Furthermore, the few white matter regions showing a positive relationship with age, in posterior and inferior portions of the brain, are located in visual, spatial, and motor areas, which show little to no decline with aging. Despite the strong evidence we have presented showing the differential timecourse of structural decline across the lifespan, these data were collected in a cross-sectional design and thus represent between-participant differences as opposed to reflecting change over time within an individual. Longitudinal data are essential to investigate whether these differences develop and progress as individuals age, and that aging of the grey and white matter are coupled over time, and follow-up is currently underway for this sample.

## Acknowledgements

This study was funded in part by grants from the National Institutes of Health R00 AG-036818, R00 AG-036848, and R01 AG-056535. We thank Giuseppe Miranda for assistance with the FreeSurfer analyses and Matt Kmiecik for comments on earlier versions of the manuscript.

## Supplemental Information

*Supplemental Figure S1. Bootstrap ratios for Surface Area FA (A) and MD (B).* Barplots show bootstrap ratios for GM surface area regions, while brain images show bootstrap ratios for WM voxels. Bootstrap ratios below an absolute value of 3 are considered non-significant and are faded in GM barplots, or green in WM voxelwise brains. Note. STS = Superior Temporal Sulcus.

*Supplemental Figure S2. Surface Area Latent Variable Projections and the Relationship of each Brain Score with Age for (A-C) FA and (D-F) MD.* Grey matter brain scores increase linearly with increased age demonstrating that data from younger adults drive the negative side of component 1, while data from older adults drive the positive side of component 1 (shown for FA in panel B and for MD in panel E). A similar pattern is observed in white matter brain scores in a nonlinear fashion (i.e., accelerates with age) (for FA in panel C and for MD in panel F). Scatter dots represent individual participants, triangles represent the mean of an age “group”, and blue color fade from dark to light blue represents younger to older adults, respectively to illustrate the age effect. Both linear and nonlinear regression lines are illustrated on the scatterplots, with the more significant fit denoted with bold solid lines, and the lower fitting line denoted by faded dashed lines. Note. YA= Younger Adults; MA = Middle-Aged Adults; OA = Older Adults; VOA = Very Old Adults.

*Supplemental Figure S3. Bootstrap ratios for GM Volume FA (A) and MD (B).* Barplots show bootstrap ratios for GM volume regions, while brain images show bootstrap ratios for WM voxels. Bootstrap ratios below an absolute value of 3 are considered non-significant and are faded in GM barplots, or green in WM voxelwise brains. Note. STS = Superior Temporal Sulcus.

*Supplemental Figure S4. GM Volume Latent Variable Projections and the Relationship of each Brain Score with Age for (A-C) FA and (D-F) MD.* Grey matter brain scores increase linearly with increased age demonstrating that data from younger adults drive the negative side of component 1, while data from older adults drive the positive side of component 1 (shown for FA in panel B and for MD in panel E). A similar pattern is observed for white matter brain scores in a nonlinear fashion (i.e., accelerates with age) (shown for FA in panel C and for MD in panel F). Scatter dots represent individual participants, triangles represent the mean of an age “group”, and blue color fade from dark to light blue represents younger to older adults, respectively to illustrate the age effect. Both linear and nonlinear regression lines are illustrated on the scatterplots, with the more significant fit denoted with bold solid lines, and the lower fitting line denoted by faded dashed lines. Note. YA= Younger Adults; MA = Middle-Aged Adults; OA = Older Adults; VOA = Very Old Adults.

