## Supplemental Figures for "Joint contributions of cortical morphometry and white matter microstructure in healthy brain aging: A partial least squares correlation analysis"

**Supplemental Information**

*Supplemental Figure S1. Bootstrap ratios for Surface Area FA (A) and MD (B).* Barplots show bootstrap ratios for GM surface area regions, while brain images show bootstrap ratios for WM voxels. Bootstrap ratios below an absolute value of 3 are considered non-significant and are faded in GM barplots, or green in WM voxelwise brains. Note. STS = Superior Temporal Sulcus.

*Supplemental Figure S2. Surface Area Latent Variable Projections and the Relationship of each Brain Score with Age for (A-C) FA and (D-F) MD*. Grey matter brain scores increase linearly with increased age demonstrating that data from younger adults drive the negative side of component 1, while data from older adults drive the positive side of component 1 (shown for FA in panel B and for MD in panel E). A similar pattern is observed in white matter brain scores in a nonlinear fashion (i.e., accelerates with age) (for FA in panel C and for MD in panel F). Scatter dots represent individual participants, triangles represent the mean of an age "group", and blue color fade from dark to light blue represents younger to older adults, respectively to illustrate the age effect. Both linear and nonlinear regression lines are illustrated on the scatterplots, with the more significant fit denoted with bold solid lines, and the lower fitting line denoted by faded dashed lines. Note. YA= Younger Adults; MA = Middle-Aged Adults; OA = Older Adults; VOA = Very Old Adults.

*Supplemental Figure S3. Bootstrap ratios for GM Volume FA (A) and MD (B).* Barplots show bootstrap ratios for GM volume regions, while brain images show bootstrap ratios for WM voxels. Bootstrap ratios below an absolute value of 3 are considered non-significant and are faded in GM barplots, or green in WM voxelwise brains. Note. STS = Superior Temporal Sulcus.

*Supplemental Figure S4. GM Volume Latent Variable Projections and the Relationship of each Brain Score with Age for (A-C) FA and (D-F) MD*. Grey matter brain scores increase linearly with increased age demonstrating that data from younger adults drive the negative side of component 1, while data from older adults drive the positive side of component 1 (shown for FA in panel B and for MD in panel E). A similar pattern is observed for white matter brain scores in a nonlinear fashion (i.e., accelerates with age) (shown for FA in panel C and for MD in panel F). Scatter dots represent individual participants, triangles represent the mean of an age "group", and blue color fade from dark to light blue represents younger to older adults, respectively to illustrate the age effect. Both linear and nonlinear regression lines are illustrated on the scatterplots, with the more significant fit denoted with bold solid lines, and the lower fitting line denoted by faded dashed lines. Note. YA= Younger Adults; MA = Middle-Aged Adults; OA = Older Adults; VOA = Very Old Adults.

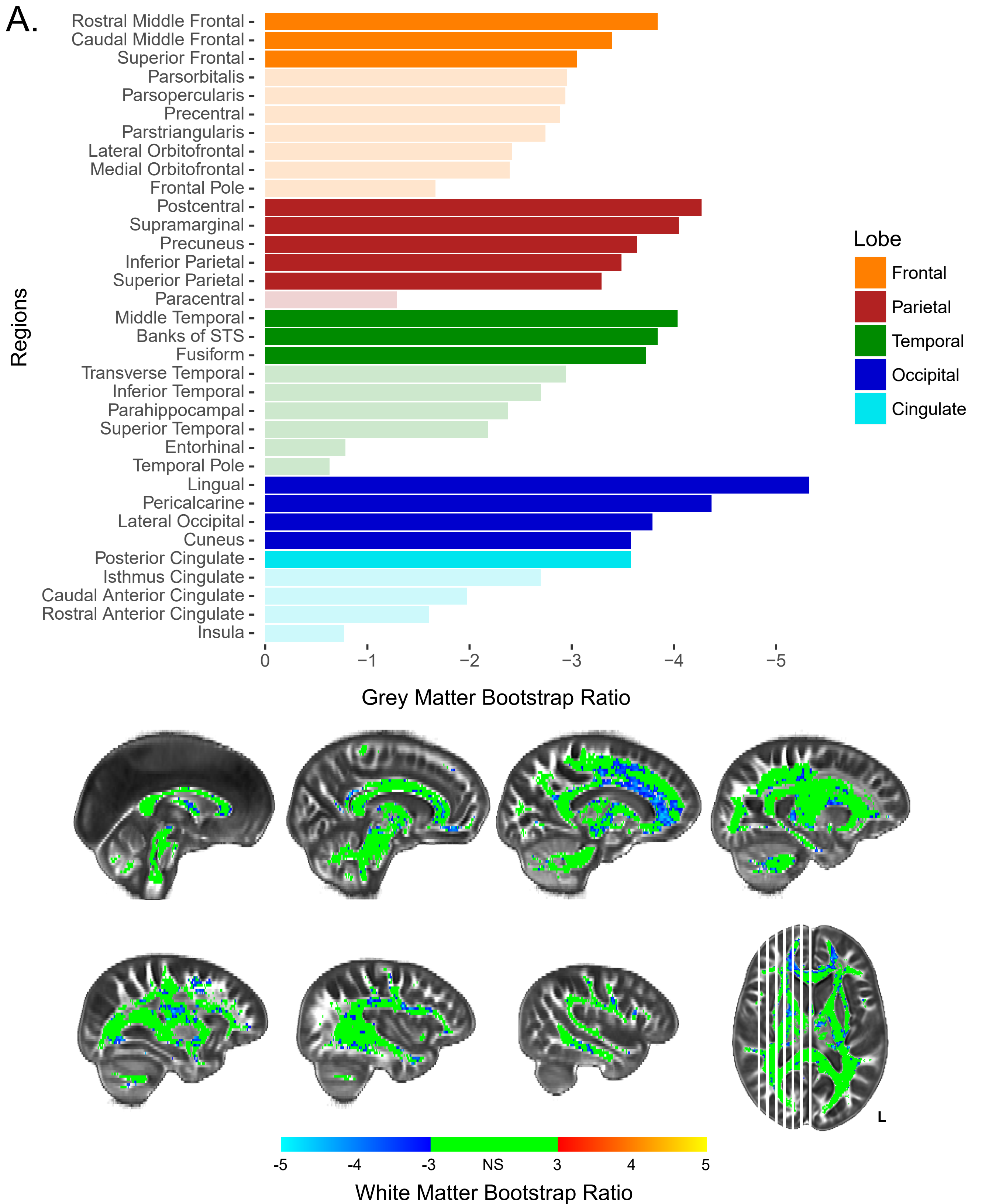
A.
Rostral Middle Frontal
Caudal Middle Frontal
Superior Frontal
Parsorbitalis
Parsopercularis
Precentral
Parstriangularis
Lateral Orbitofrontal
Medial Orbitofrontal
Frontal Pole
Postcentral
Supramarginal
Precuneus
Inferior Parietal
Superior Parietal
Paracentral
Middle Temporal
Banks of STS
Fusiform
Transverse Temporal
Inferior Temporal
Parahippocampal
Superior Temporal
Entorhinal
Temporal Pole
Lingual
Pericalcarine
Lateral Occipital
Cuneus
Posterior Cingulate
Isthmus Cingulate
Caudal Anterior Cingulate
Rostral Anterior Cingulate
Insula
Regions
0
−1
−2
−3
−4
−5
Grey Matter Bootstrap Ratio
Lobe
Frontal
Parietal
Temporal
Occipital
Cingulate
L
-5
-4
-3
NS
3
4
5
White Matter Bootstrap Ratio

B.

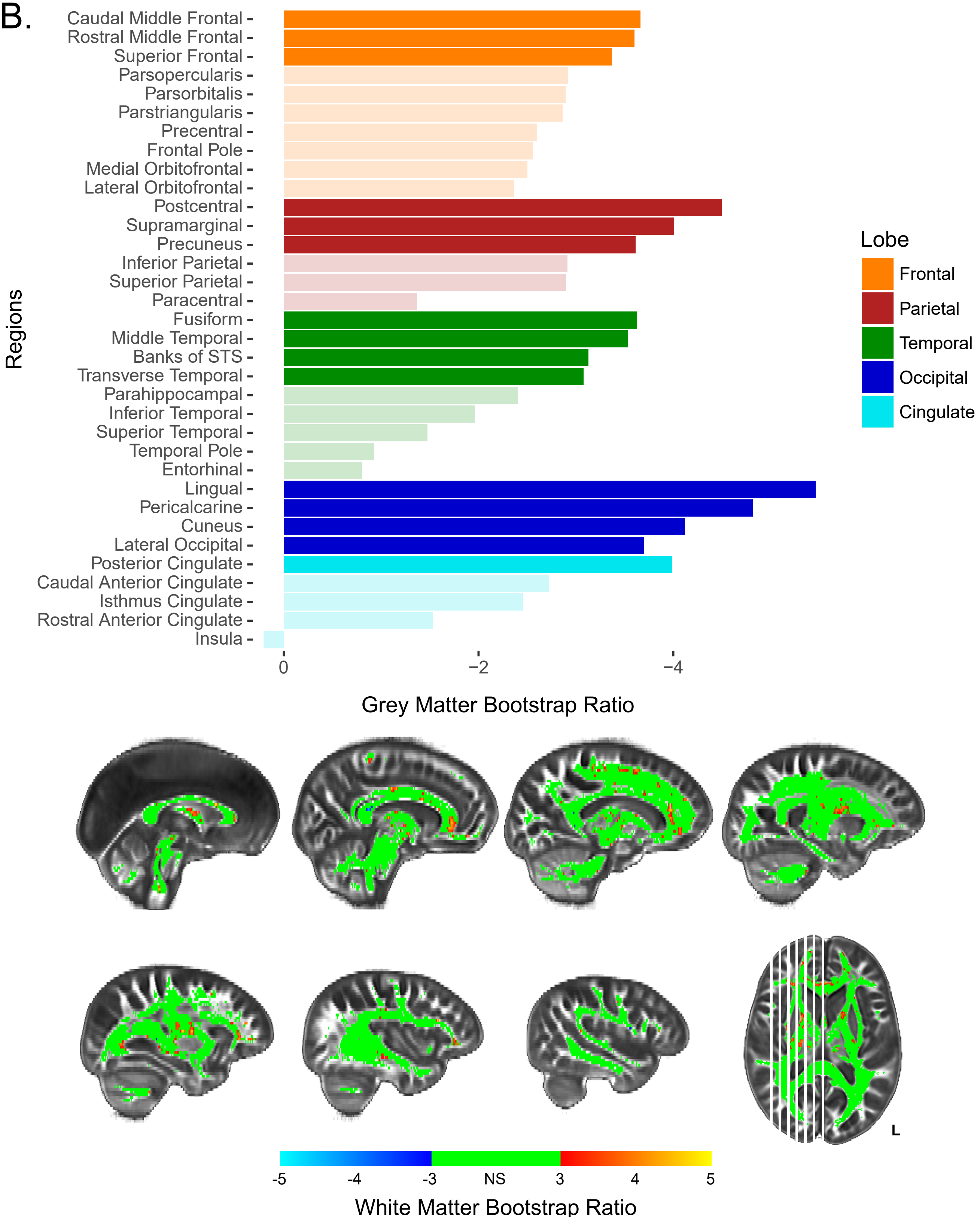
Caudal Middle Frontal
Rostral Middle Frontal
Superior Frontal
Parsopercularis
Parsorbitalis
Parstriangularis
Precentral
Frontal Pole
Medial Orbitofrontal
Lateral Orbitofrontal
Postcentral
Supramarginal
Precuneus
Inferior Parietal
Superior Parietal
Paracentral
Fusiform
Middle Temporal
Banks of STS
Transverse Temporal
Parahippocampal
Inferior Temporal
Superior Temporal
Temporal Pole
Entorhinal
Lingual
Pericalcarine
Cuneus
Lateral Occipital
Posterior Cingulate
Caudal Anterior Cingulate
Isthmus Cingulate
Rostral Anterior Cingulate
Insula
Regions
0
−2
−4
Grey Matter Bootstrap Ratio
Lobe
Frontal
Parietal
Temporal
Occipital
Cingulate
L
-5
-4
-3
NS
3
4
5
White Matter Bootstrap Ratio

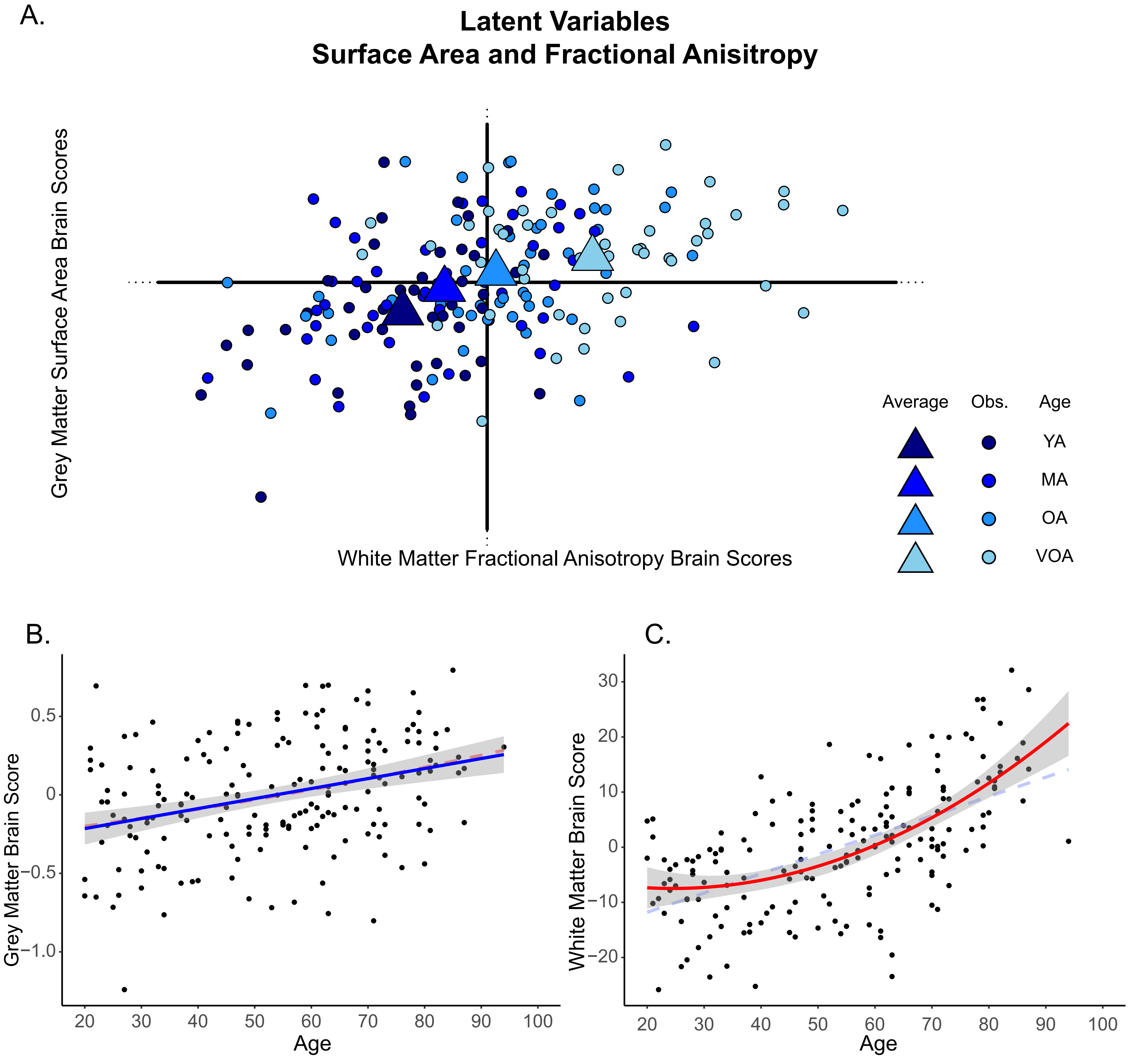

A.
Latent Variables
Surface Area and Fractional Anisitropy
Grey Matter Surface Area Brain Scores
White Matter Fractional Anisotropy Brain Scores
Average
Obs.
Age
YA
MA
OA
VOA
B.
Grey Matter Brain Score
0.5
0
−0.5
−1.0
20
30
40
50
60
70
80
90
100
Age
C.
White Matter Brain Score
30
20
10
0
−10
−20
20
30
40
50
60
70
80
90
100
Age

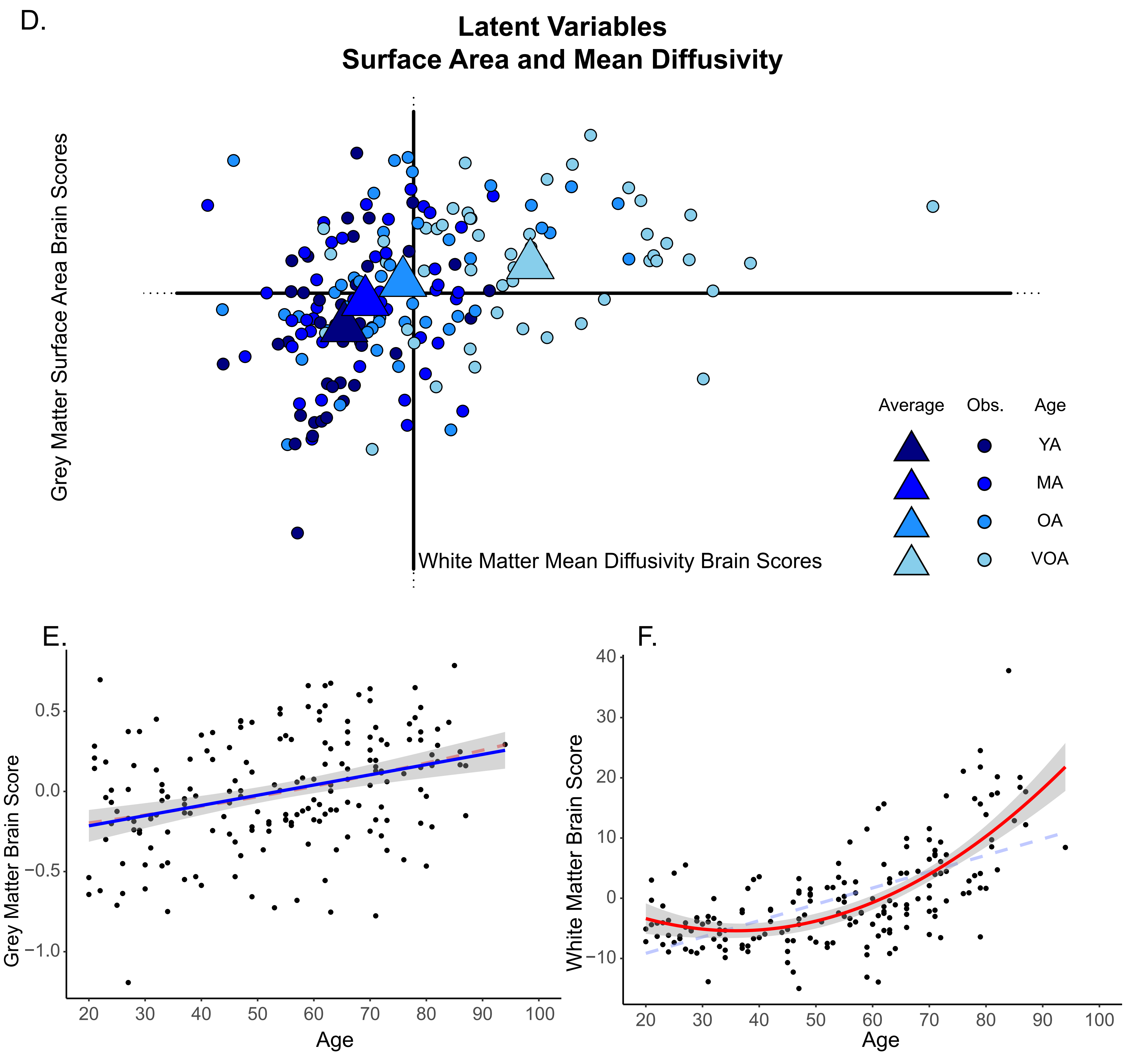

D.
Latent Variables
Surface Area and Mean Diffusivity
Grey Matter Surface Area Brain Scores
White Matter Mean Diffusivity Brain Scores
Average
Obs.
Age
YA
MA
OA
VOA
E.
Grey Matter Brain Score
0.5
0.0
−0.5
−1.0
20
30
40
50
60
70
80
90
100
Age
F.
White Matter Brain Score
40
30
20
10
0
−10
20
30
40
50
60
70
80
90
100
Age

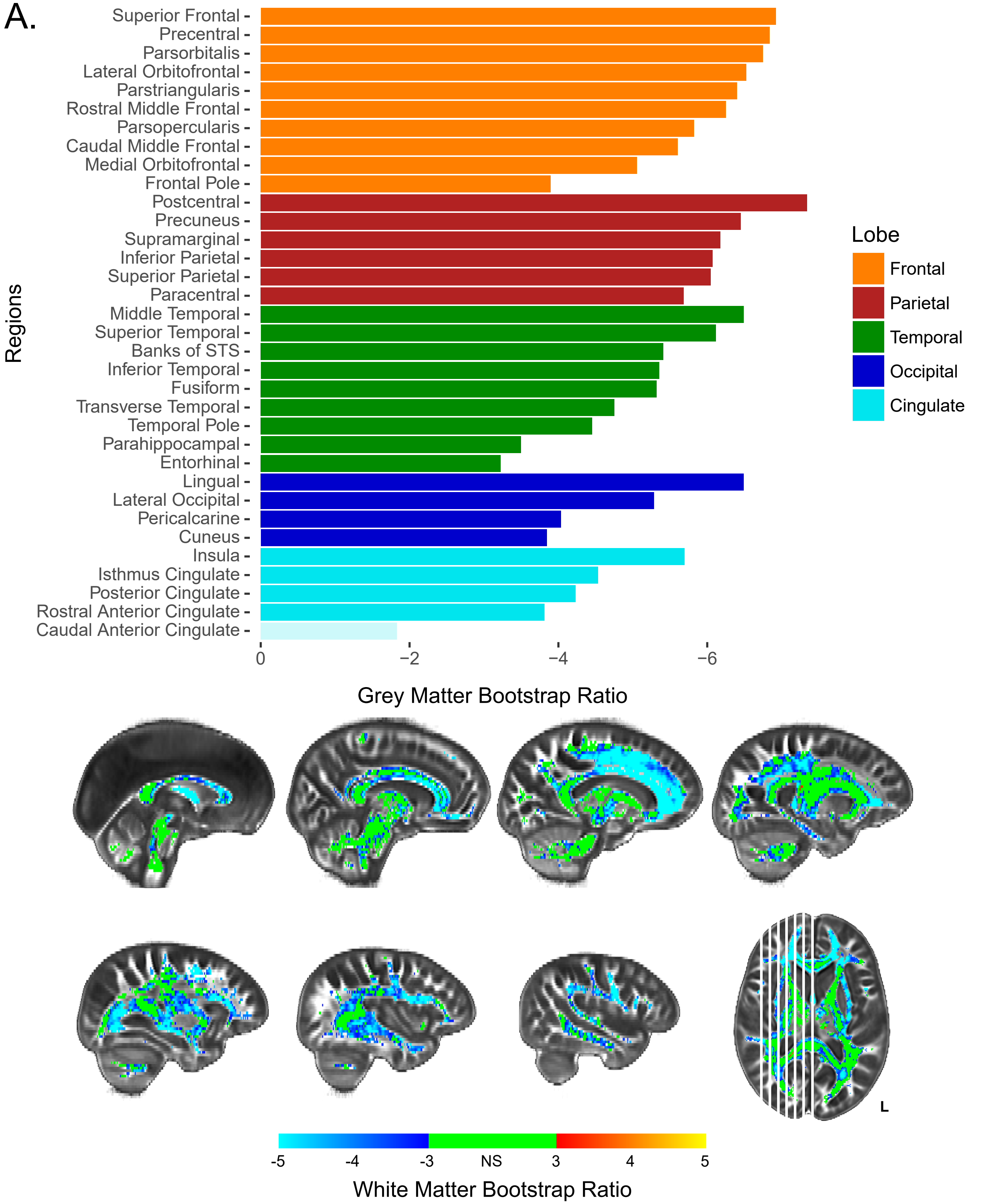
A.
Superior Frontal
Precentral
Parsorbitalis
Lateral Orbitofrontal
Parstriangularis
Rostral Middle Frontal
Parsopercularis
Caudal Middle Frontal
Medial Orbitofrontal
Frontal Pole
Postcentral
Precuneus
Supramarginal
Inferior Parietal
Superior Parietal
Paracentral
Middle Temporal
Superior Temporal
Banks of STS
Inferior Temporal
Fusiform
Transverse Temporal
Temporal Pole
Parahippocampal
Entorhinal
Lingual
Lateral Occipital
Pericalcarine
Cuneus
Insula
Isthmus Cingulate
Posterior Cingulate
Rostral Anterior Cingulate
Caudal Anterior Cingulate
Regions
0
−2
−4
−6
Grey Matter Bootstrap Ratio
Lobe
Frontal
Parietal
Temporal
Occipital
Cingulate
L
-5
-4
-3
NS
3
4
5
White Matter Bootstrap Ratio

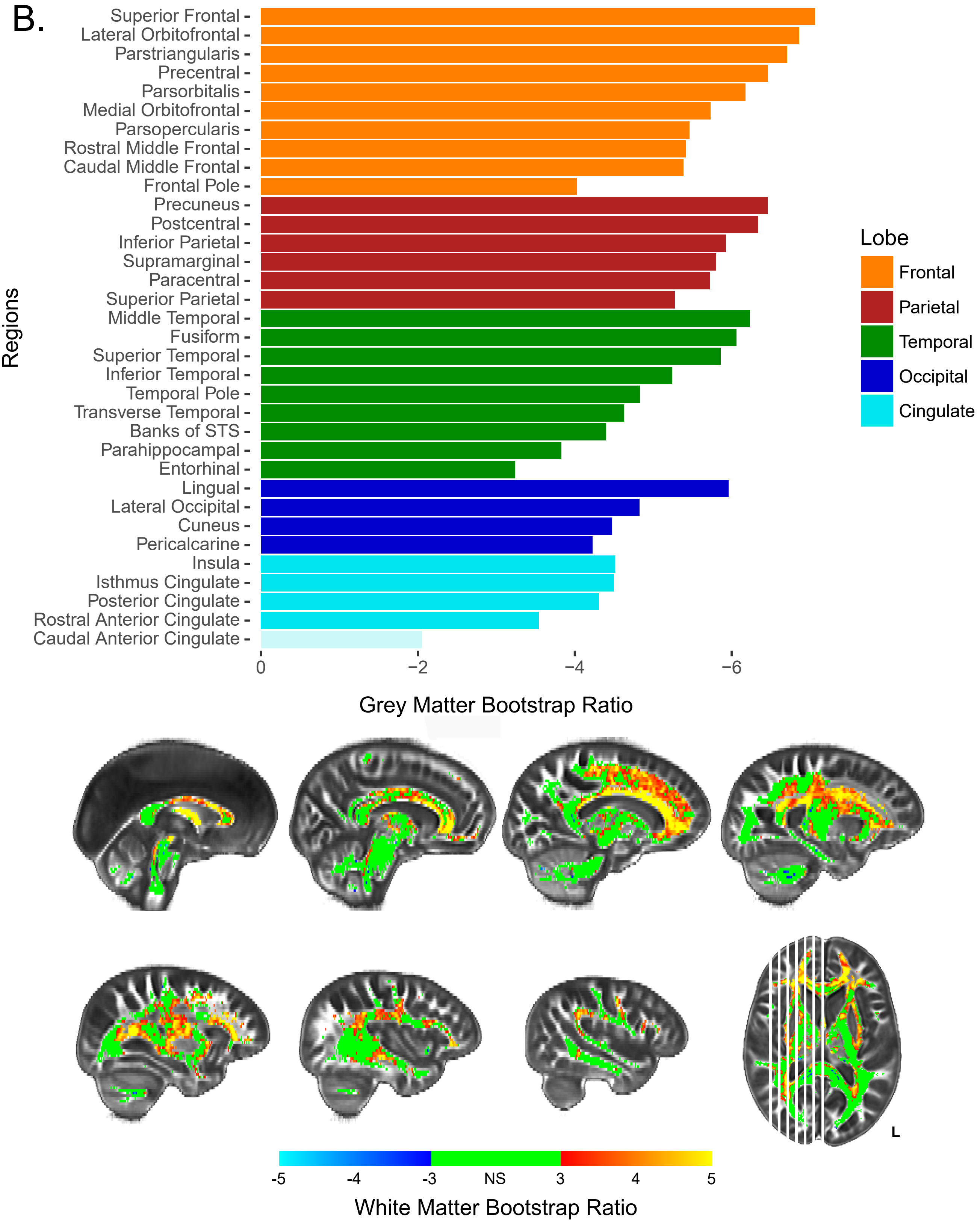
B.
Superior Frontal
Lateral Orbitofrontal
Parstriangularis
Precentral
Parsorbitalis
Medial Orbitofrontal
Parsopercularis
Rostral Middle Frontal
Caudal Middle Frontal
Frontal Pole
Precuneus
Postcentral
Inferior Parietal
Supramarginal
Paracentral
Superior Parietal
Middle Temporal
Fusiform
Superior Temporal
Inferior Temporal
Temporal Pole
Transverse Temporal
Banks of STS
Parahippocampal
Entorhinal
Lingual
Lateral Occipital
Cuneus
Pericalcarine
Insula
Isthmus Cingulate
Posterior Cingulate
Rostral Anterior Cingulate
Caudal Anterior Cingulate
Regions
0
−2
−4
−6
Grey Matter Bootstrap Ratio
Lobe
Frontal
Parietal
Temporal
Occipital
Cingulate
L
-5
-4
-3
NS
3
4
5
White Matter Bootstrap Ratio

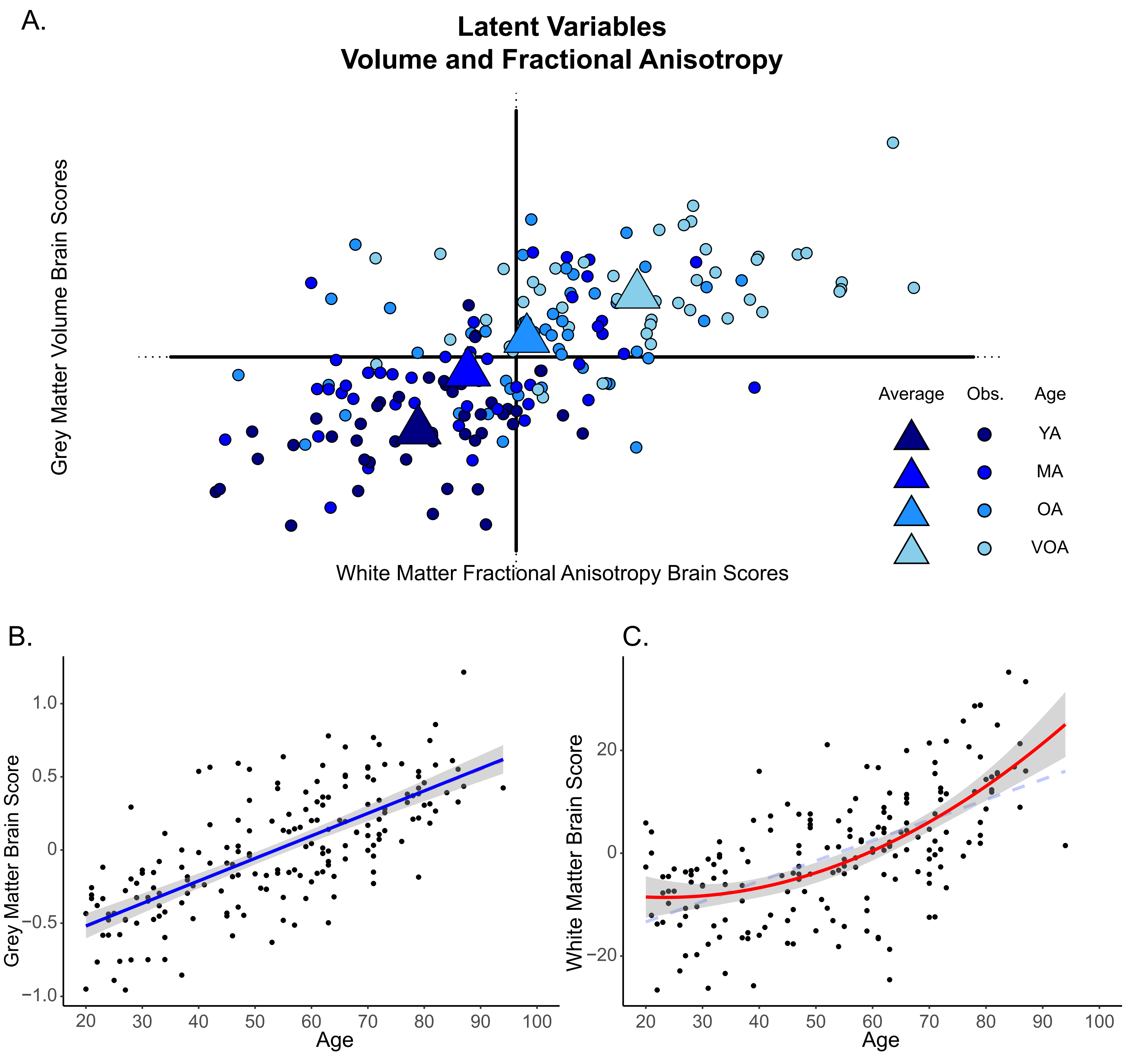
A.
Latent Variables
Volume and Fractional Anisotropy
Grey Matter Volume Brain Scores
White Matter Fractional Anisotropy Brain Scores
Average
Obs.
Age
YA
MA
OA
VOA
B.
Grey Matter Brain Score
1.0
0.5
0
−0.5
−1.0
20
30
40
50
60
70
80
90
100
Age
C.
White Matter Brain Score
20
0
−20
20
30
40
50
60
70
80
90
100
Age

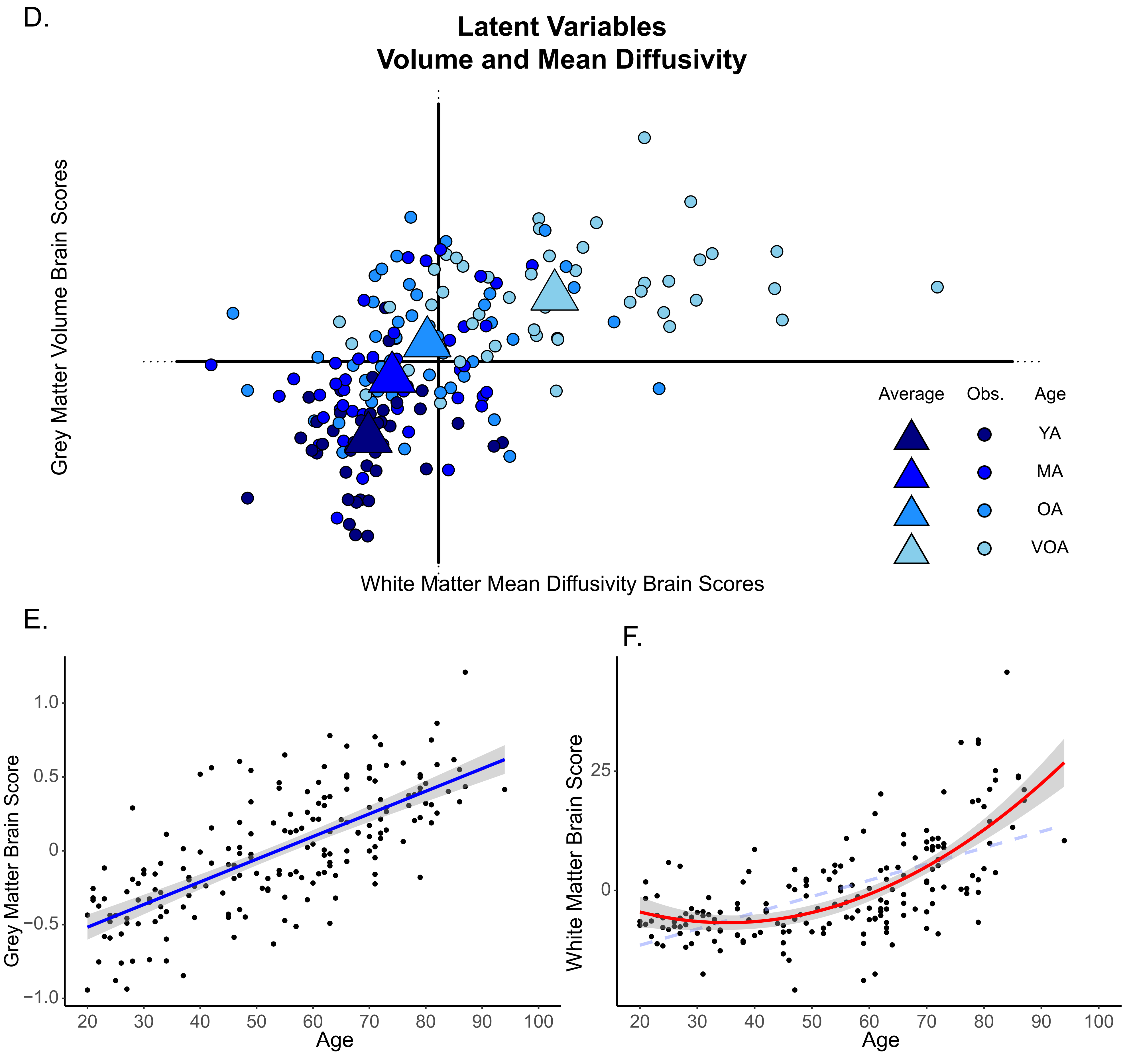

D.
Latent Variables
Volume and Mean Diffusivity
Grey Matter Volume Brain Scores
White Matter Mean Diffusivity Brain Scores
Average
Obs.
Age
YA
MA
OA
VOA
E.
1.0
0.5
0
−0.5
−1.0
Grey Matter Brain Score
20
30
40
50
60
70
80
90
100
Age
F.
25
0
White Matter Brain Score
20
30
40
50
60
70
80
90
100
Age
